# A robust Platform for Integrative Spatial Multi-omics Analysis to Map Immune Responses to SARS-CoV-2 infection in Lung Tissues

**DOI:** 10.1101/2023.02.19.529128

**Authors:** Xiao Tan, Laura F. Grice, Minh Tran, Onkar Mulay, James Monkman, Tony Blick, Tuan Vo, Ana Clara Simões Flórido Almeida, Jarbas da Silva Motta Junior, Karen Fernandes de Moura, Cleber Machado-Souza, Paulo Souza-Fonseca-Guimaraes, Cristina Pellegrino Baena, Lucia de Noronha, Fernanda Simoes Fortes Guimaraes, Hung N. Luu, Tingsheng Drennon, Stephen Williams, Jacob Stern, Cedric Uytingco, Liuliu Pan, Andy Nam, Caroline Cooper, Kirsty Short, Gabrielle T. Belz, Fernando Souza-Fonseca-Guimaraes, Arutha Kulasinghe, Quan Nguyen

## Abstract

The SARS-CoV-2 (COVID-19) virus has caused a devastating global pandemic of respiratory illness. To understand viral pathogenesis, methods are available for studying dissociated cells in blood, nasal samples, bronchoalveolar lavage fluid, and similar, but a robust platform for deep tissue characterisation of molecular and cellular responses to virus infection in the lungs is still lacking. We developed an innovative spatial multi-omics platform to investigate COVID-19-infected lung tissues. Five tissue-profiling technologies were combined by a novel computational mapping methodology to comprehensively characterise and compare the transcriptome and targeted proteome of virus infected and uninfected tissues. By integrating spatial transcriptomics data (Visium, GeoMx and RNAScope) and proteomics data (CODEX and PhenoImager HT) at different cellular resolutions across lung tissues, we found strong evidence for macrophage infiltration and defined the broader microenvironment surrounding these cells. By comparing infected and uninfected samples, we found an increase in cytokine signalling and interferon responses at different sites in the lung and showed spatial heterogeneity in the expression level of these pathways. These data demonstrate that integrative spatial multi-omics platforms can be broadly applied to gain a deeper understanding of viral effects on cellular environments at the site of infection and to increase our understanding of the impact of SARS-CoV-2 on the lungs.

## Introduction

### The need for tissue profiling at sites of infection

The COVID-19 pandemic has led to over 618 million total infections and more than 6.5 million deaths globally (https://coronavirus.jhu.edu/, as of Oct 2022). While the lungs are the main site affected by the severe acute respiratory syndrome coronavirus 2 (SARS-CoV-2), the high degree of heterogeneity in SARS-CoV-2 infected lung cells at different lung locations is not well understood (1). Whilst vaccinations have helped reduce disease severity and viral spread, the occurrence of viral-induced fibrosis, which can lead to complications in the lung and other organs, is still common (2, 3). Cytokines such as interleukin-6 (IL-6) and transforming growth factor-β have been shown to contribute to pulmonary fibrosis (4) suggesting that understanding of the interactions between immune and other cells in the infected tissue are required to prevent infection-induced tissue damage. However, it is still largely unclear how inflammatory immune cells become activated and induce damage in organs such as the lungs. Moreover, in preparation to fight future pandemics, robust clinical research platforms are needed to rapidly determine the responsible pathogens, characterise the host immune response and define the pathogenic mechanisms at both the cellular and molecular levels directly from patient tissues (5, 6).

### Multi-omic assessment provides a comprehensive view of cellular and molecular pathogenesis of the virus

During the COVID-19 pandemic, genomics and bioinformatics have proven to be important public health tools in applications ranging from screening to tracking variants (5, 7, 8). Multi-omics enables the systematic understanding of virus pathogenesis (8). The successful application of an mRNA vaccine as the main vaccine solution globally further suggests the need for more transcriptomics research at the scale of either the targeted gene/gene-set or transcriptome-wide levels. Furthermore, adding proteomics, phospho-proteomics and ubiquitination data can reveal the role of post-transcriptional regulation and virus-host interactions at protein level (9, 10). Application of single-cell technologies enables viral effects to be studied across organs and reveals cell-specific immune responses (11, 12). In addition, multi-omics integration increases statistical power to analyse small patient cohorts, resulting in reliable patient classification models using molecular markers (13)

### A spatial multi-omics approach to experimentally and computationally study infected lung tissues

So far, molecular studies of COVID-19 infection have mainly been limited to studying dissociated tissues, blood samples, nasal swabs, and bronchoalveolar lavage fluid, but viral activity and spread within the lungs have not been fully investigated (12, 14). Spatial-omics technologies are an area of rapid development resulting in the swift evolution of methods to profile RNA or protein with spatial context within infected tissues, and increasing tools for analysis. Spatial analysis shows the upregulation of interferon (IFN) type I response (15) and upregulation at the alveolar regions of IFN-α, IFN-γ and oxidative phosphorylation pathways (11). However, each spatial technology presents unique benefits and limitations with regards to detection sensitivity and resolution, background signal, and the number of molecules that can be measured (16). In this context, using independent technologies in parallel is required to examine all possible genes and cell types and to cross-validate observations at protein and RNA levels.

Here, we report one of the first studies to implement multi-modal spatial technologies to study COVID-19 response. We applied five spatial-omics technologies to the same tissue biopsies to provide complementary information about RNA and protein expression at different genetic resolutions (ranging from as many as >20,000 genes to a small set of six curated proteins). Spatial context was retained for all technologies. We spatially profiled blood vessels, lung alveolar type II pneumocytes (T2), immune regions, and epithelial tissue regions, comparing lung samples from patients with and without SARS-CoV-2, and those with high and low levels of SARS-CoV-2 viral mRNA signal. Importantly, we developed a computational pipeline to enable spatial integration of protein and RNA data across tissue sections. The integration and comparisons revealed cell types and molecular pathways of potential importance to virus pathogenesis.

## Results

### Building a multi-omics tissue atlas of COVID-19

To comprehensively understand changes in spatial organisation of cells in lung tissue sections infected by SARS-CoV-2 at the RNA, protein and cell type levels, we applied five different spatial technologies to serial sections of tissue microarrays (TMAs) generated from patient biopsies (**Fig 1**). The key sample set in this study included four COVID-19-infected samples (referred to as LN1 to LN4), which were each measured using all five technologies. We also included additional uninfected samples for certain technologies, in total measuring six uninfected samples for CODEX, five for GeoMx, and nine for PhenoImager HT (**Fig 1**).

**Figure 1.**
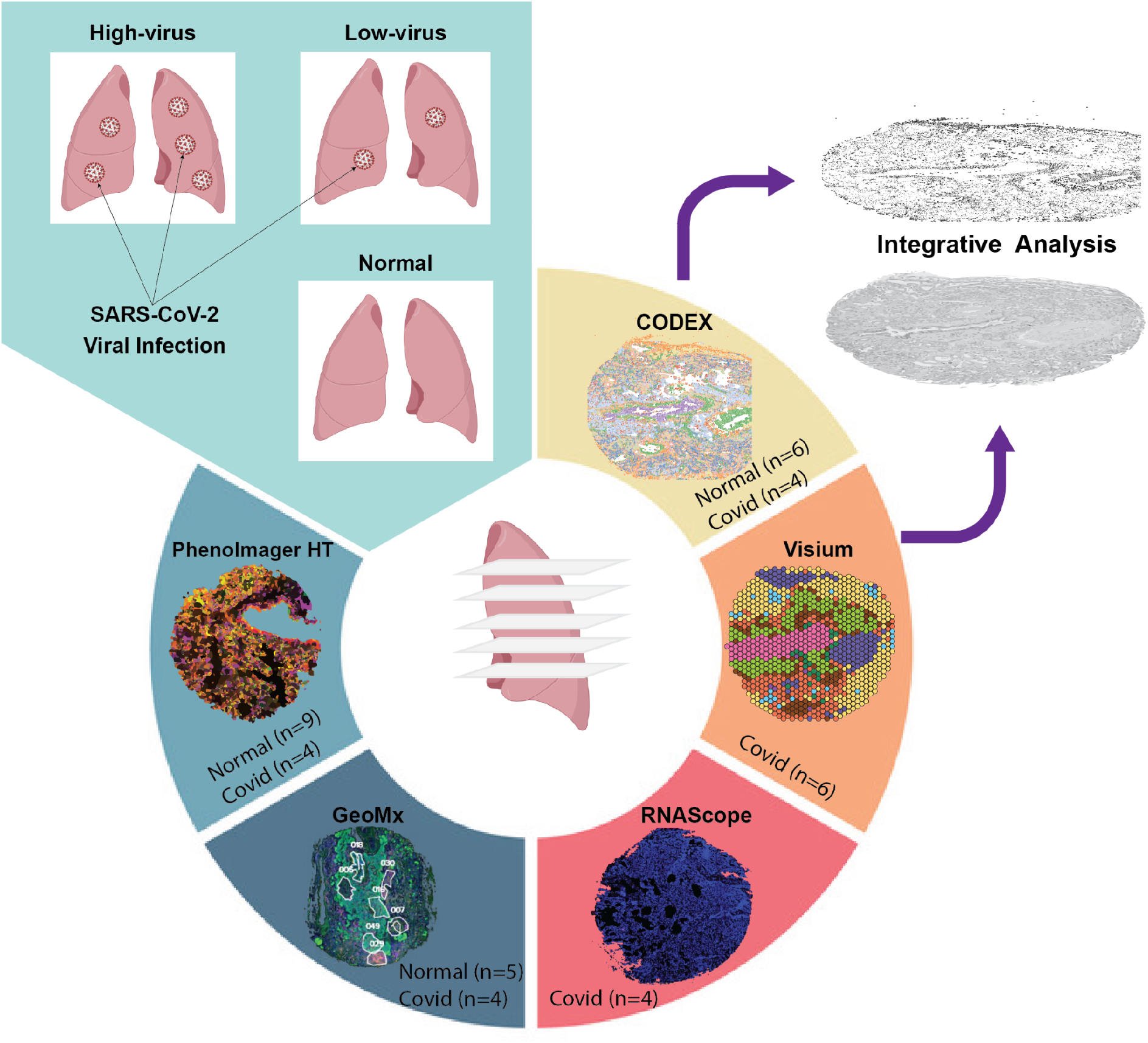
A spatial multi-omics approach to mapping molecular and cellular responses to SARS-CoV-2 infection in the lungs. Five complementary spatial RNA and protein technologies were applied to the same lung core biopsies. Adjacent tissue sections (CODEX and Visium) were mapped for data integration across modalities. In addition to the four core biopsy samples captured with all five data types, we profiled additional samples totalling 13 for Phenoimager HT, nine for GeoMX, four for RNAScope, six for Visium and 10 for CODEX. Together, the multimodal dataset enables deep comparison between COVID-19 and normal healthy lung tissues and viral high vs low viral samples. Representative tissue images from a single COVID-19-infected sample, LN2, are shown in each segment.

To detect the presence of SARS-CoV-2 mRNA signalling the four core biopsy samples (**Fig S1a**), we first applied RNAScope to target the SARS-CoV-2 spike mRNA, followed by quantification and localisation of the virus by our STRISH pipeline (17). This analysis classified samples as two viral mRNA signal high- and two viral signal low-samples and also defined the spatial location of the virus within lung tissues (**Fig S1b**). We used this classification in subsequent analyses comparing molecular differences between samples based on differing viral mRNA signal. To compare responses to infection at the transcriptional level in an unbiased manner, we produced Visium data from the four core biopsy samples, which after pre-processing captured non-zero gene expression data for 17,680 genes, with the average number of genes per Visium spot ranging from 564 to 2,698 across a total of 129 to 969 spots per biopsy. To uncover gene expression changes across different selected morphological regions, we applied GeoMx whole transcriptome atlas (WTA) technology for five uninfected and four COVID-19 patients (total 49 regions of interest; ROIs). Approximately 18,000 genes were measured across four morphological categories annotated by a pathologist as Blood vessel, T2 (Type II pneumocytes), Immune, and Epithelial regions. While RNAScope, Visium and GeoMx modalities captured cellular RNA content, we also measured protein expression using CODEX for six uninfected and four COVID-19-infected samples. A total of 36 proteins were captured for each sample. Furthermore, high-resolution multiplexed protein imaging via PhenoImager HT was performed to detect six key immune markers, namelyCD56 (NK cells), IFI27, CD15 (neutrophils), CD66b (granulocytes), CD8 (cytotoxic T cells), and CD3 (total T cells). Together, the five complementary technologies form a unique spatial multi-omics dataset to unbiasedly study transcriptional and proteomic changes in COVID-19 responses in lung tissues and between patients in an unbiased manner (**Fig 1**).

### Spatial distribution and changes in cell type composition of COVID-19-infected and uninfected lung tissues

Computational clustering, annotation, community detection and deconvolution methods were applied to the five data types to compare cell types across the four COVID-19-infected lung samples. Histological features were captured using H&E imaging from Visium data (**Fig S1a**), corresponding to samples with high (LN1 and LN3) and low (LN2 and LN4) viral signal (**Fig S1b**). Spatial cell type distribution is shown in **Fig S1** for the five data types, including Visium (**Fig S1c; Fig S4**), CODEX (**Fig S1d**), PhenoImager HT (**Fig S1e-f**), and GeoMX (**Fig S1g-i)**

We first sought to map the major lung cell types in spatial context within the Visium data using label transfer (18) based on an intermediate-level cell type annotation from a published scRNASeq COVID-19 atlas (19). We identified 11 dominant cell types, three of which were shared across all four patient samples (airway epithelial cells, fibroblasts, and plasma cells) (**Fig S4b**). Molecular gene signatures for two key cell types, namely airway epithelial cells and smooth muscle cells, highlight two anatomical features in one tissue section (**Fig 2b-c**). These molecularly-defined structures matched closely with anatomical structures visible in the underlying H&E image (**Fig S1a**). These structures were not detected in other samples using either morphological or molecular data, suggesting that the cell type detection in Visium data is highly tissue-specific. From GeoMx data, we uncovered high heterogeneity in cell type composition across different lung regions including blood vessels, epithelium, type II pneumocytes-rich regions, and immune-rich regions (**Figure S1i**).

**Figure 2.**
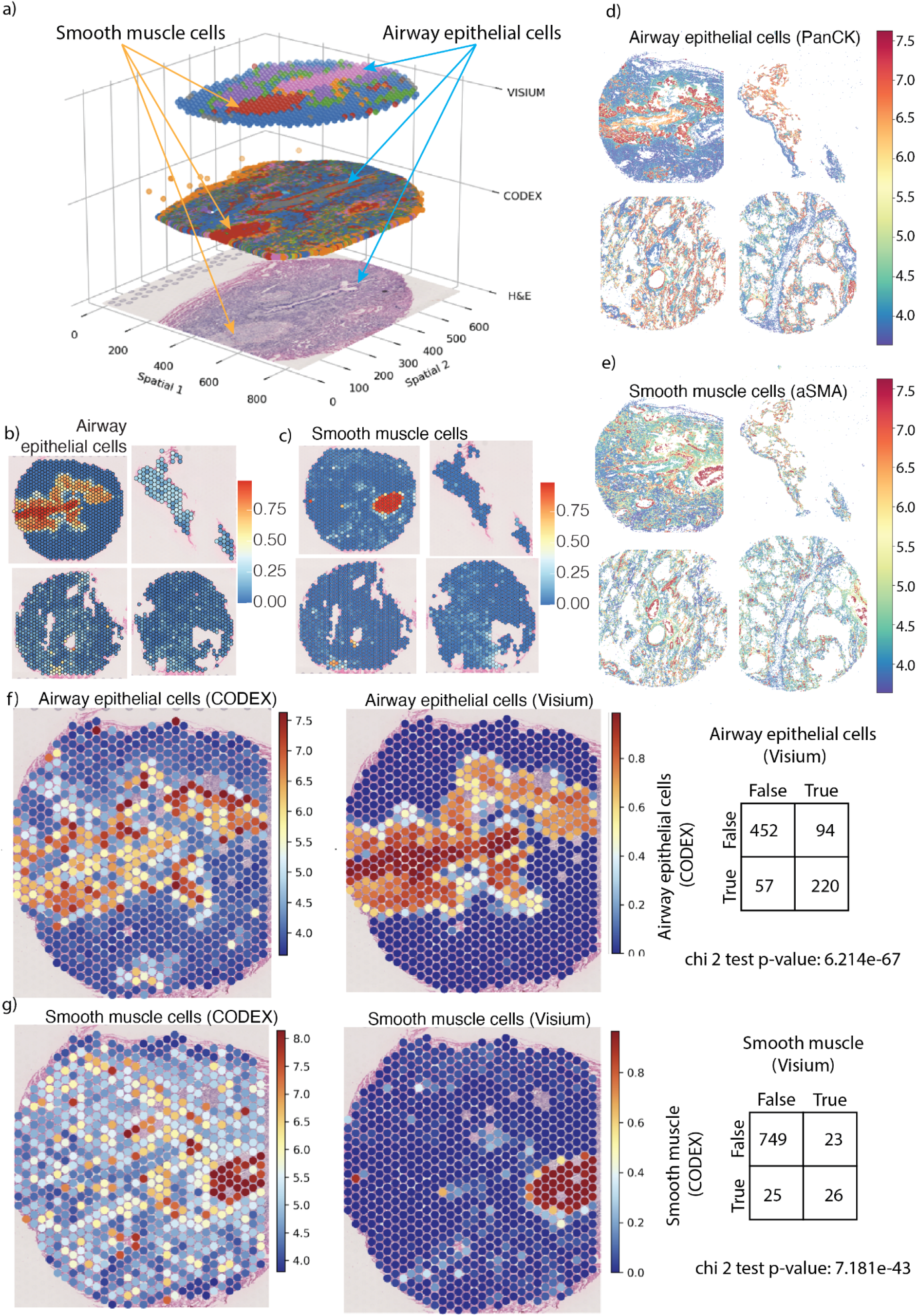
Computational transfer of molecular expression from CODEX protein single-cell data to Visium RNA spot-level data. **a**) Integrative analysis based on automated mapping of CODEX protein data to Visium RNA data and the Visium H&E tissue image. The three layers show, from top to bottom, Visium clustering results, CODEX clusters, and the corresponding H&E tissue image captured in the Visium experiment. Data for sample LN2 is shown. (**b-c**) Visium cell type classification showing distribution of cell type scores for epithelial (b) and smooth muscle (c) cells. Scores were determined using the label transfer method (18) with reference data from a publicly-available COVID-19 lung atlas (19). (**d-e**) CODEX protein expression for epithelial marker PanCK (d) and smooth muscle cell marker αSMA (e). (**f**) Data transfer of CODEX protein marker expression to Visium spots. Left - PanCK expression from CODEX data mapped within each Visium spot region. Middle - label transfer scores from **Panel b** for visual comparison with CODEX protein expression. Right - comparison between transferred CODEX protein data and Visium RNA-based cell type labelling. The contingency table shows the consistency in spots classified as airway epithelial cells in the Visium data and spots that contain the transferred positive PanCK signal from CODEX data, with Chi-square test (p value < 0.05). The positive PanCK Z-score signal was used to classify positive vs negative PanCK spots (refer to the method section). (**g**) As in **Panel f**, but comparing CODEX expression of smooth muscle marker αSMA (left) and Visium smooth muscle annotations (middle).

We performed further cell type composition comparisons between COVID-19-infected and uninfected samples using PhenoImager HT (**Fig S1 e-f**) and GeoMx deconvolution data (**Fig S1h-i**). Consistently, both PhenoImager HT and GeoMx data suggest a significant increase in NK cells in COVID-19-infected samples compared to uninfected samples (**Fig S1f-g**). The GeoMx data also suggested that the infected samples had more CD4 T cells than uninfected samples (**Fig S1g**,**i**); these cell types were not captured by the PhenoImager HT protein panel. While CD8 T cells were significantly higher in COVID-19 samples based on PhenoImager HT data, this difference was not apparent in the GeoMx data (although the median gene expression value for relevant markers was indeed higher in COVID-19 samples) (**Fig S1g-i**). CODEX RNA data also showed higher expression of the CD8 T cell activation marker, *CD107a* in infected patients (**Fig S7**). The increase in CD8 T cell numbers has been previously shown in COVID-19 studies using single cell data analysis (12). Similarly, the PhenoImager HT data showed higher neutrophils (marked as CD66^+^) in COVID-19 samples, but this trend was not clear in the deconvolution results from GeoMx data (**Fig S1g-i**). Exacerbated neutrophil response in COVID-19 patients has been previously observed (20). Overall, GeoMx data appeared to under detect the difference in CD8 T cells and neutrophils. This may be due to the fact that the CD8 and CD66 protein markers were used directly as definitive classifiers for CD8 T cells and neutrophils for CODEX data, while these two cell types were inferred by combinatorial signatures of multiple genes in GeoMx data.

### Differential gene activity underlying high- and low-viral COVID-19 infection

We detected a total of 2,132 genes in the Visium data whose expression differed between samples with high- and low-mRNA viral signal. This total included 115 genes that were upregulated in the high-viral samples and 2,017 genes that were upregulated in the low-viral sample (**Fig S4c-d**). Interestingly, these gene sets could potentially be used as signatures for classifying tissues with high vs low virus signal, as we observed similar enrichment of the low-viral gene signature in two independent Visium samples from a lung tissue microarray which were shown by RNAScope to have a low viral mRNA signal(**Fig S6**). Further, we found that the suite of genes upregulated in the high-virus samples was enriched in pathways associated with immune function. For instance, the differentially expressed gene list was enriched for immune-related gene ontology (GO) terms relating to interferon response (**Fig S4e**), antigen presentation, chemokine signalling, and regulation of viral genome replication (**Fig S4f**). This pathway level finding is not only consistent with previous reports (11, 12, 15, 20), but also extends the associated gene list to include potential new candidate genes which were not detectable in by previous analysis but were identified using the transcriptome-wide Visium assay.

Using GeoMx data, we performed an unsupervised approach to identify those of the 18,000 captured genes that were differentially expressed

### Site-specific comparisons of gene expression in the lung

with GeoMx region labels, we compared different tissue types (Bronchial Epithelium, Blood Vessel, between COVID-19 vs uninfected samples. The top genes upregulated in COVID-19 infected samples compared to uninfected samples included an MHC class I gene *B2M*, a cytokine macrophage migration inhibitory factor *CD74*, inflammatory gene *LAP3*, and genes controlling alveolar surface tension like *SFTPA2* and *SFTPB* (**Fig S2e**). We also compared the differentially expressed genes to three curated lists of COVID-19-relevant genes, namely those associated with interferon responses, apoptosis responses, and prognostic markers(**Fig S2a-c**). Several apoptosis markers like *CASP4, BAK1*, and *BAX* were significantly upregulated in the infected samples (**Fig S2a**). A range of some, but not all, known interferon markers were also upregulated in our data, including *IFI6, IFIT1, LY6E*, and *IFR7* (**Fig S2b**). Among the 11 potential prognostic markers investigated, we found four of them to be highly expressed in COVID-19 samples.

Top upregulated genes from a further unsupervised comparison between high vs low viral mRNA signal samples included those related to interferon response (e.g. *IFIT3, IRF1 and IFI6*) and to chemokine inflammatory responses (e.g. *CCL8* and *CXCL11*) (**Fig S2d**).

T2 pneumocyte region) separately in infected and uninfected samples. This analysis identified 495 genes that were consistently upregulated in the infected samples (**Fig S2f**). These genes are significantly enriched in key pathways such as signalling by interleukins, influenza infection, antigen processing-cross presentation, viral mRNA translation (**Fig S2g, h**).

When focussing specifically on T2 (type II pneumocyte)-containing ROIs, we identified 281 differentially expressed genes between COVID-19 infected and uninfected samples. This list included cytokine markers upregulated in infected samples, such as *CXCL2, CXCL8, CXCL9, CXCL16, CCL4L2, IL6* and *IL18BP* (**Fig S2f**).

### Integration of multi-omic imaging data cross-validates tissue types and spatial expression

We developed an innovative computational approach to automatically integrate spatial multi-omics data based on first cross-mapping tissue images and transferring assayed molecular data between samples. We here automatically cross-mapped Visium and CODEX data via image registration and spatial coordinate transformation (**Fig 2a**). This mapping allowed us to transfer all 36 CODEX proteins to the Visium H&E image, which already provided measurements of ∼17,680 genes. This generated an integrated dataset anchored by histological H&E tissue images from Visium that contained both multiplexed protein and spatial transcriptomics data. The three levels of information in the integrated dataset (tissue morphological information, RNA counts and protein expression) enabled powerful analyses combining pathological annotation with transcriptome-wide gene expression and high-plex protein data.

To evaluate the mapping results, we examined the identification of airway epithelial cells and smooth muscle cells in the LN2 samples, as these two cell types both appeared in well-defined histopathological regions that could be used as a reliable ground-truth reference. Visual inspection of the spatial distribution of Visium cell type scores (**Fig 2b-c)** revealed clearly-defined tissue zones corresponding to the airway epithelium and smooth muscle tissue, respectively. The CODEX single-cell resolution expression of markers for epithelial cells (PanCK) and smooth muscle (αSMA) clearly suggested that the locations of the smooth muscle and the airway in this data modality (**Fig 2d-e**) are consistent with the Visium cell type annotation. However, the CODEX signals exhibited greater spread across the tissue sections.

For quantitative comparison across data modalities, we used the integrated data to map CODEX protein signals back to coordinates corresponding to each Visium spot (**Fig 2f-g**). This mapping approach allowed us to directly compare Visium spots that are classified as epithelial cells or smooth muscle cells with the same pseudo-spots in the CODEX data that are positive or negative for the PanCK or αSMA markers, respectively. Assessment of the resulting contingency tables and statistical testing showed that the numbers of true positive and true negative spots in classifying the two cell types were significantly higher than random cell type assignment (**Fig 2f-g**). This observation suggests that by integration of independent spatial transcriptomics and spatial proteomics data, we can cross-validate cell types across different -omics technologies.

### Macrophage distribution differs with infection status

Here we used Visium, CODEX, and GeoMx data to further investigate host responses to infection by mapping macrophage distribution and spatial cytokine expression across the tissue sections. We compared samples with high vs low viral signal (LN1 and LN3 vs LN2 and LN4 respectively), and those with vs without COVID-19 infection (**Fig 2f-g**).

Both Visium and CODEX data indicated a strong macrophage presence across the tissues. However, macrophage abundance was clearly enriched in the high viral mRNA signal samples (LN1 and LN3) in the Visium data compared to the low viral signal samples (**Fig 3a**). We observed that macrophages tended to cluster around the airway region in sample LN2 **(Fig 2b, d)**, based on both overall transcriptional signatures in Visium data (**Fig 3a**) or the expression of key macrophage protein markers CD163, CD68, CD14 and CD11b in CODEX data (**Fig 3b**). Comparison of COVID-19-infected and uninfected samples did not indicate a significant difference in macrophage distribution, but did highlight that macrophages distribute surrounding airway structures (**Fig 3b, Fig 4d**).

**Figure 3.**
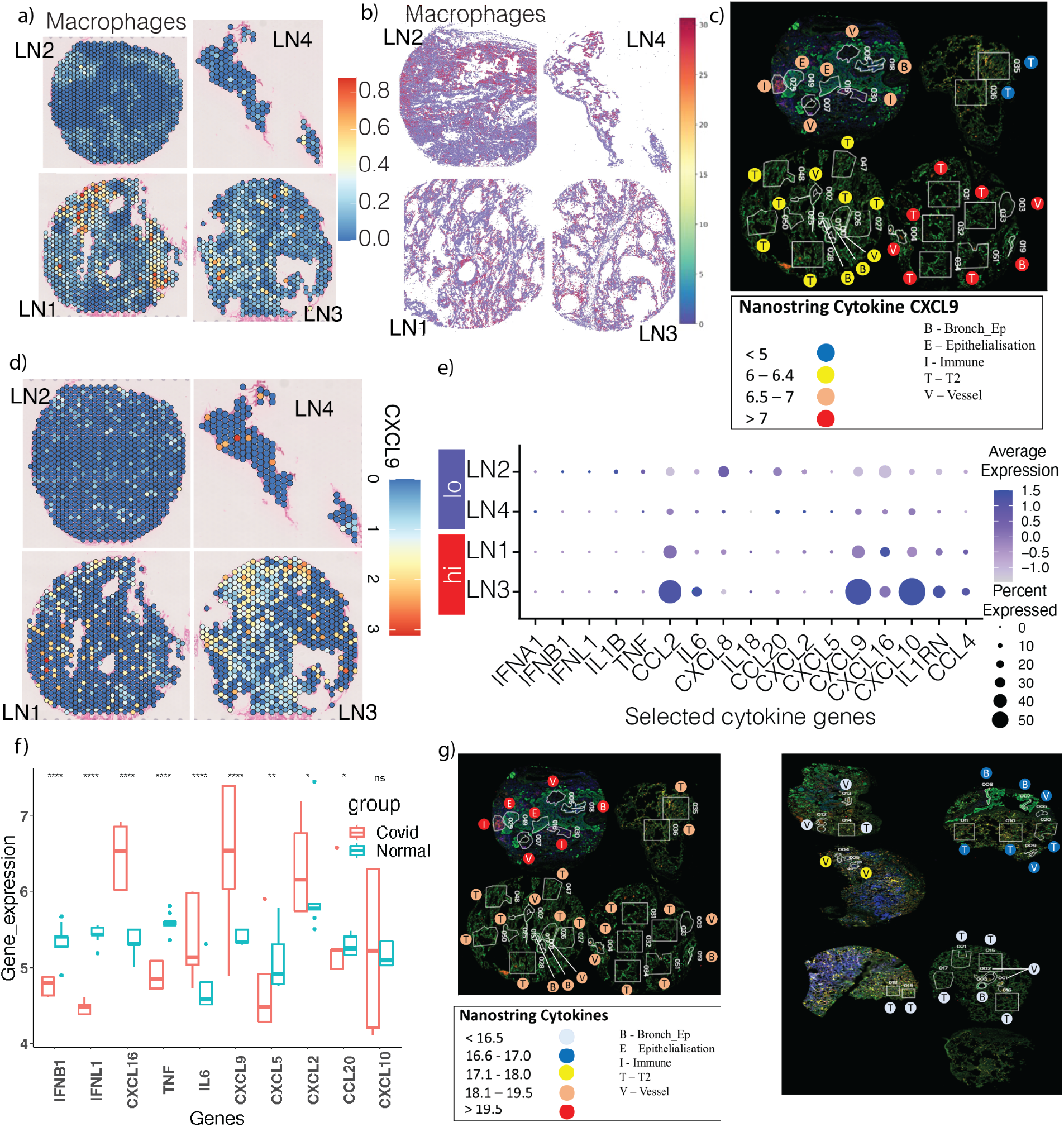
Spatial multi-omics assessment of macrophage and cytokine responses to SARS-CoV-2 infection. (**a**) Mapping of macrophages using Visium data. The colour scale indicates the proportion of each spot predicted to contain macrophages. Infected lung tissues from four COVID-19 patients are shown. LN1 and LN3 are samples with a high viral mRNA signal, while LN2 and LN4 have a low viral mRNA signal. (**b**) CODEX protein expression of three macrophage markers, showing the sum of the normalised CODEX signal for CD163, CD68, CD14, and CD11b. (**c**) Expression of exemplar cytokine gene CXCL9 in GeoMx data. Log Counts Per Million (logCPM) transformed fitted values from a glm model are shown for the four COVID-19 samples. Each ROI (white boxes) is annotated with a circle indicating gene expression (colour) and tissue annotation (letter). (**d**) Spatial expression pattern of the CXCL9 cytokine in Visium data for four tissues, two with low viral signal and two with high viral signal. (**e**) Expression of a curated set of cytokine genes in Visium data. Dot colour indicates gene expression levels and dot size indicates the percentage of Visium spots in each sample expressing the gene of interest. (**f**) Statistical differences in gene expression for cytokines (IFNB1, IFNL1, CXCL16, TNF, IL6, CXCL9, CXCL5, CXCL2, CCL20, and CXCL10) in GeoMx data between COVID-19-infected and uninfected tissues. (**g**) Average log CPM-transformed fitted values from a glm model comparing COVID-19-infected and uninfected tissues for cytokine CXCL2, CXCL9, and CXCL16. Each ROI (white boxes) is annotated with a circle indicating gene expression (colour) and tissue annotation (letter).

**Figure 4.**
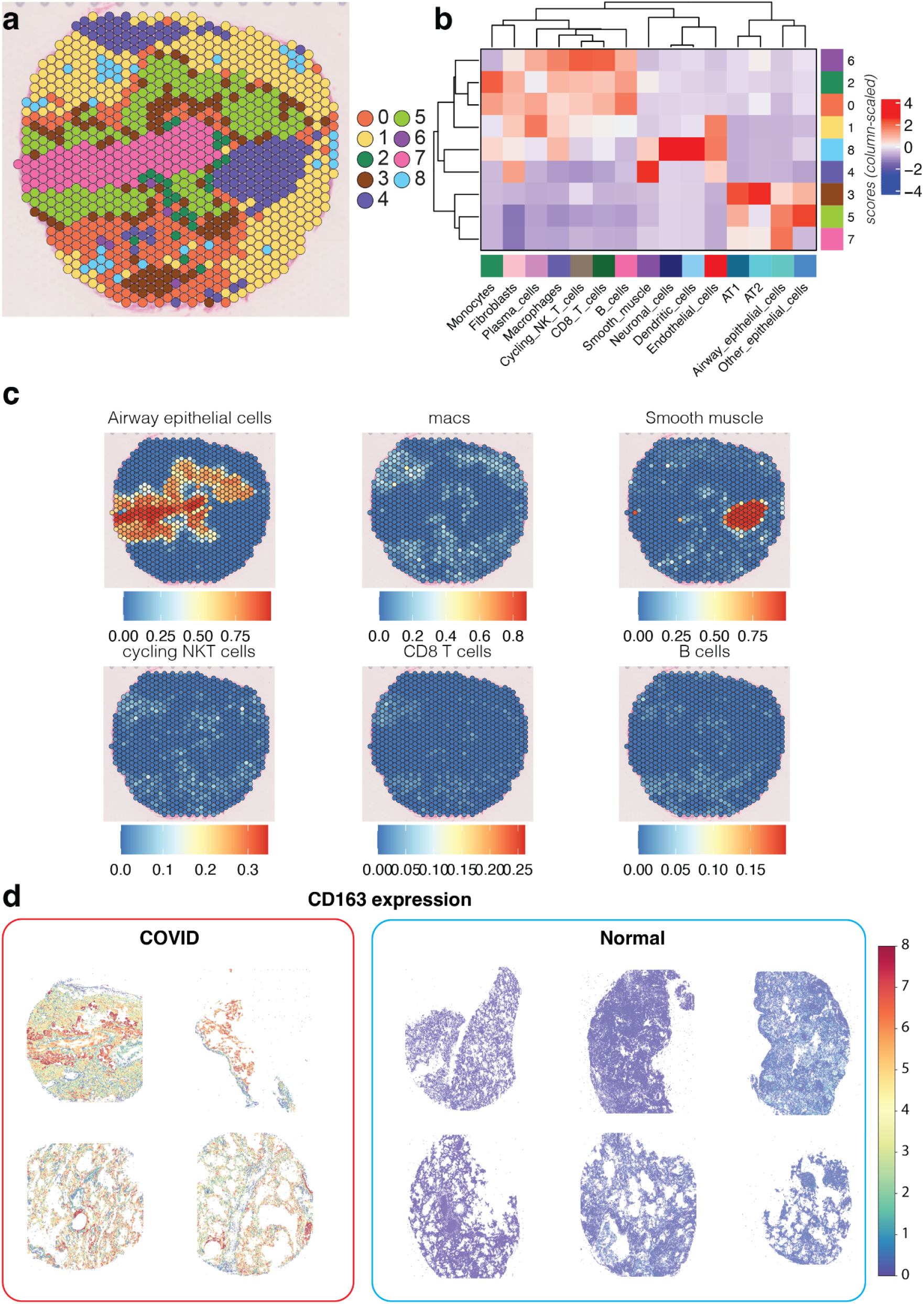
Cell type predictions in a COVID-19 infected sample (LN2 patient) by using both Visium and Phenoimager HT data. (**a**) Results of clustering on integrated data (clustering resolution of 0.6), showing patient sample LN2 only. (**b**) Heatmap indicating the average label transfer score of each cell type in each predicted cell cluster. Values are scaled by column (cell type). (**c**) Label transfer scores for key cell types shown in B plotted to the tissue. (**d**) Spatial expression of macrophage markers in COVID-19 infected and normal lung tissue consistently shows a much stronger presence of macrophages in the COVID-19 infected samples.

We also used Visium data to investigate macrophage distribution within their cellular communities. Here, we performed unbiased clustering to group Visium capture spots (**Fig 4a**). Due to the non-single-cell resolution of Visium data, these clusters represent communities of multiple cell types; we therefore determined the populations of cell types present within each cluster (**Fig 4b**). We thereby found communities where macrophages colocalised with CD8 T cells, B cells, NK cells and plasma cells (**Fig 4b**; clusters 6, 2 and 0). Qualitative inspection of our label transfer spot annotation results for sample LN2, without consideration of the underlying clusters, further revealed that macrophages tended to be situated next to AT1 cells, overlapped NK cells, B cells and CD8 T cells, and sat near AT1 and AT2 cells. However, fibroblasts and macrophages did not overlap and instead were localised to different tissue compartments distinct from one another (**Fig 4c; Fig S5**).

### Spatial multi-omics mapping of cytokine expression changes at RNA and protein levels

We then compared cytokine expression levels across data modalities, because cytokine storms have been particularly associated with severe or lethal clinical outcomes in COVID-19 patients (21). We focussed on a curated list of cytokines previously shown to be active in COVID-19-infected in postmortem lung samples (22, 23) (**Fig 3c, f-g** for GeoMx, **Fig 3d-e** for Visium). Overall, we found a significant increase in cytokine expression in COVID-19-infected samples compared to uninfected samples (**Fig 3f-g**) and again in infected samples with high viral signal compared to those with lower viral signal (**Fig 3e**). In particular, we observed the upregulation of cytokines *CXCL16, IL6* and *CXCL2* in an infection context (**Fig 3f**). However, this trend is not universal; both GeoMx and Visium data suggested that several other cytokines, including *CXCL20, CXCL5, TNF, IFNB1*, and *IFNL1* did not increase in infected samples or in samples with a higher virus signal (**Fig 3e-f**).

We next specifically analysed the spatial distribution of a single cytokine gene, *CXCL9*, which showed the greatest difference between samples with low and high viral signal (**Fig 3e**) and infected and uninfected samples (**Fig 3f**) in both Visium and GeoMx data, respectively. CXCL9 is an IFN-γ-induced ligand of CXCR3 expressed by macrophages (24) which has been shown to be upregulated during COVID-19 infection in humans and mice (25), a finding consistent with our results. GeoMx and Visium data both indicate that *CXCL9* expression is highest in high-viral mRNA signal patient sample LN3 (**Fig 3c-d**).

At the protein modality, the CODEX data was tested for statistically significant differences in protein expression between COVID-19 infected and uninfected patients. We found higher expression of CD45, CD163 (monocyte/macrophage marker), and CD107a (cytotoxic CD8 T cell marker) in COVID-19 samples compared to uninfected ones (**Fig S7**), an observation consistent with the results from the GeoMx RNA data.

## Discussion

The global COVID-19 pandemic has proven the essential roles of genomics and advanced multi-omics analyses as public health tools and discovery platforms for RNA vaccines, immune responses, and pathogenic mechanisms of infection (5, 6). A key feature of COVID-19 infection is the diverse molecular and cellular mechanisms underlying its highly heterogeneous initiation, severity and progression between patients. A comprehensive and multi-dimensional view of human cells responding to virus infection and spread in the lung is needed to increase readiness for future pandemics and to better understand the effects of Long COVID (12, 26). Multi-omics analysis, making use of interactome, proteome, and transcriptome data, has shown the potential to advance our understanding and help identify candidate drug targets against COVID-19 (27). One key area where we need to glean more information is the spatial characteristics of the lung tissues as the major site for virus localisation, as it is still unclear whether peripheral blood or nasal swab samples accurately reflect viral loads and effects within the lung tissue itself. Here we present an in-depth spatial multi-omics analysis of COVID-19-infected lung tissues, reconstructing a spatial map of immune responses to COVID-19 infection across lung biopsies from multiple patients.

Method development to integrate spatial multi-omics data remains at an early stage of development (26). Here, we developed an automated computational approach to map spatial data to cross-validate multimodal information about spatial gene/protein expression, cell/neighbourhood distribution and anatomical annotation on an H&E image. This approach uses an image registration method to transfer labels (i.e. annotations and/or expression information) from adjacent or nearby tissue sections to the same common coordinate for the integrative analysis (28). For tissue sections in the same block, but are further away, the integration is more challenging and less accurate, but the tissue landmarks are still mappable. We here successfully mapped CODEX protein signals to Visium RNA data. This integration confirmed cell types predicted from spatial transcriptomics data, as shown by the high correlation of CODEX protein signals and Visium RNA signals in airway epithelial cells and smooth muscle cells. We believe that this integration method is also a powerful tool for other projects, because it allows for the flexible integration of multiple data modalities measured from separate sections into a single common framework for downstream analysis.

By using Visium data and GeoMx data at bulk/pseudo-bulk levels, we identified suites of genes (n = 2,132 genes for Visium and n = 241 genes for GeoMx) associated with high-or low-virus mRNA samples. The list of genes upregulated in high-virus samples was enriched for immune-related functions including interferon response, chemokine signalling and antigen presentation. We validated our result in a replicate of an additional sample from a separate Visium data cohort. These samples showed a strong transcriptional signature for genes associated with low viral samples, which is consistent with their SARS-CoV-2 spike mRNA expression levels as determined by RNAScope. This suggests that we have identified a potential transcriptional signature that allows sample viral mRNA signal to be inferred without directly detecting the virus.

Using orthogonal Visium, CODEX and GeoMx data, we consistently found the presence of macrophages in COVID-19-infected samples, particularly those with a high viral mRNA signal. We also detected a wide range of cytokines likely to have been produced by these macrophages as part of the inflammatory responses to COVID-19 infection; again, cytokine expression was enriched in COVID-19 infected samples, more so in the samples with high viral signal. Independent datasets from GeoMx, CODEX and PhenoImager HT exhibited a shared patterns of increased NK cells and T cells in COVID-19 samples compared to uninfected samples.

Interestingly, our community analysis suggested the colocalisation of macrophages with CD8 T cells, B cells, NK cells and plasma cells. This spatial distribution pattern is consistent with a report studying bronchoalveolar lavage fluid, which showed that a rich presence of T cells and monocytes induced cytokine release from macrophages (29). Macrophages are known to contribute to cytokine storm (30). From our spatial mapping of macrophages within their immune environment, we suggest potential spatial signatures in immune responses by the host lung tissues to COVID-19 virus infection, and that this molecular signature may correlate with tissue damage and cytokine storm events.

Overall, we have developed a platform for integrative analysis of spatial multi-omics data to study the molecular and cellular mechanisms of COVID-19 infection processes at sites of infection. We expect that this platform and similar approaches will contribute to enhancing readiness for future global infection pandemics, allowing comprehensive and timely understanding of pathogenic factors and host responses.

## Methods

### Ethical approval and collection of COVID-19-infected lung sample

Autopsy and biopsy materials were obtained from the Pontificia Universidade Catolica do Parana National Commission for Research Ethics under the following ethics approval numbers: protocol number 3.944.734/2020 (for the COVID-19 group), and 2.550.445/2018 (for the control groups). The control groups did not have COVID-19 infections, but did exhibit non-viral comorbidities. All methods were carried out following relevant guidelines and regulations. Families permitted the post-mortem biopsy of COVID-19 samples and conventional autopsy for the cases in the control group. The study was approved under the University of Queensland Human Research Ethics Committee ratification.

### Multispectral multiplex IHC [PhenoImager HT]

A multispectral MOTIF panel was developed (Akoya Biosciences, US) and staining was performed on a Leica Bond RX (Leica biosystems, US) at the Walter and Eliza Hall Institute (WEHI) histology core (Melbourne, Australia). Briefly, antibodies were validated and antigen retrieval conditions optimised by DAB staining of Tonsil and Lung tissue, prior to testing in a multiplex panel. Antibodies were then tested for stable staining patterns in several panel orders. The final inal panel was 1: IFI27, abcam #ab224133, 1:30, Opal 520; 2: CD15, Biolegend #301902, 1:200, Opal 570; 3: CD66b, Biolegend #305102, 1:200, Opal 620; 4: CD8, CST #70306, 1:200, Opal 690; 5: CD56, CST #99746, 1:50; Opal 480; 6: CD3, Dako #A0452, 1:500, TSA-Dig-780. Whole slide scans were performed on Vectra PhenoImager HT (Akoya Biosciences, US) by WEHI histology core, and images were spectrally unmixed in InForm (Akoya Biosciences, US) using MOTIF spectral libraries.

### CODEX dataset generation and pre-processing

CODEX (Akoya Biosciences, US) staining was performed by Enable Medicine, US, as described previously (15). Briefly, FFPE tissue samples were mounted on 20×20mm poly-lysine treated coverslips. After antigen retrieval, 190uL of antibody solution was added to the coverslips, which were incubated for 3hrs at RT in a humidity chamber. This was followed by several cycles of washing and fixation steps prior to cyclical visualisation. Antibody targets included aSMA, CD107a, CD117, CD11b, CD11c, CD14, CD141, CD15, CD163, CD183, CD197, CD20, CD21, CD31, CD34, CD38, CD3e, CD4, CD45, CD45RA, CD45RO, CD56, CD68, CD8, FoxP3, GATA3, GranzymeB, HLA-DR, Ki67, PanCK, PGP9.5, Podoplanin, RORgammaT, Siglec8, and Vimentin. For a more detailed protocol, see the reference (31).

Coverslips were imaged on an inverted fluorescence microscope (Keyence BX-810) using a Plan Apo 20x 0.75 NA objective (Nikon). The Codex imaging cycles were performed using a Codex Instrument (Akoya Biosciences) to image each barcoded antibody with corresponding barcoded fluorophore. Large regions were broken up into tiled subregions, and five z-stack slices were imaged with a step size of 1.5 μm.

Images were deconvolved and pre-processed using an image pre-processing pipeline (Enable Medicine, US). Briefly, background signal was removed from the image by using a computationally aligned blank acquisition cycle as a reference channel. Then, image deconvolution was performed for each biomarker image z-stack, and the best focus was chosen using an extended depth of field algorithm. Finally, the individual tiles were aligned and stitched together, and all channels were stacked. Image analysis was performed by Enable Medicine (US). Nuclear cell segmentation was performed using *DeepCell*, followed by segmentation dilation (32). Cellular protein expression levels were computed from the mean fluorophore intensity for each biomarker, and .fcs data were exported for downstream analysis.

The raw protein expression intensity matrices were first filtered by quantile. Cells with total counts lower than 0.05 quantile or higher than 0.95 quantile were discarded to remove the outliers. Filtered expression matrices were subtracted by *Blank* and *Empty* channels then log-transformed. Cytoskeletal *Vimentin* and neuronal *PGP9*.*5* were also discarded from further analysis.

### CODEX dataset cell type Identification

Cell type identification was performed using the Python package *Scanpy (33)*. PCA analysis was firstly applied to pre-processed CODEX datasets. Top 15 PCs were then used to calculate the neighbouring graph (*n*_*neighbour*=5). Finally, the graph-based clustering method Louvain was performed to reveal the cell types.

### Pseudo-bulk DE analysis for CODEX data to compare COVID-19-infected samples vs uninfected

Four COVID-19 samples were selected to perform differentially expressed (DE) analysis against six non-COVID-19 samples. The single cell level pre-processed protein expression was first aggregated by tissue samples using *aggregateAcrossCells()* function in *Scater* (34) package and then normalised by library size, using sample-specific normalisation factors calculated by the function *calcNormFactors()* in *edgeR* package (35). Each tissue sample was treated as pseudo-bulk data to fit in a gene-wise linear model *glmQLFit()*, which estimates quasi-likelihood dispersions between species and samples. We then implemented empirical Bayes quasi-likelihood F-tests from *glmQLFTest()* function to identify the DE genes (FDR < 0.05, log_2_-fold change > 1 or log_2_-fold change < -1). The DE genes were visualised in heatmaps by using *pheatmap* package and in volcano plots by using *EnhancedVolcano* package (https://github.com/kevinblighe/EnhancedVolcano).

### Visium data processing and analysis methods

Visium spatial transcriptomics data was gathered from a total of 2,615 capture spots across the four patient biopsies, with the median number of reads per spot ranging from 393 (LN4) to 5103 (LN2) (**Fig S1b**). Raw sequencing reads were processed and mapped to against the *Homo sapiens* genome GRCh38-2020A (CellRanger v1.2.2, 10x Genomics); the resulting filtered count matrices were used for downstream analyses. Preliminary data processing was performed to remove spots with fewer than 100 reads or genes, or with a mitochondrial or ribosomal read count >50%. Individual tissue samples were demultiplexed from their original tissue array and re-integrated using canonical correlation analysis (CCA) based on the top 2000 variable genes (18). Data normalisation was performed using scran v1.14.6 (36) and data scaling, dimensionality reduction (based on the top 15 principal components), clustering and sub-clustering, data integration, and marker prediction were performed in Seurat v4.0.0 (37)(18, 37). Clustering was tested using a range of resolution values from 0 to 1.2 and the highest average stable resolution value was selected for each sample using the SC3 measure from Clustree (38). Unbiased clustering of spots resulted in the identification of nine clusters, five of which were present (i.e. n_spots_ ≥ 5) in all tissues and two of which were tissue-specific (cluster 6 in LN3 and cluster 8 in LN2). Clusters 4 and 7 recapitulate clear anatomical regions in sample LN2, representing smooth muscle and airway epithelial cells, respectively (**Fig S3a**). Label transfer was used to annotate spots with their major cell type representative, using the intermediate-level annotation from Melms et al.’s scRNASeq COVID-19 atlas (19) (**Fig S3b**). This analysis predicted the presence of epithelial cells (airway epithelial cells, alveolar epithelial type I and type II cells (AT1 and AT2)), immune cells (CD4+ T cells, cycling NK/T cells, macrophages, plasma cells), smooth muscle, fibroblasts, endothelial cells, neuronal cells. The four dominant cell types were fibroblasts (n_spots_ = 796), plasma cells (n_spots_ = 469), alveolar type II cells (n_spots_ = 796), and airway epithelial cells (n_spots_ = 420).

As each Visium spot is composed of multiple cells (approximately 1 - 9 cells per spot), we visualised the label transfer scores for each individual cell type in turn (**Fig S4**). We observed that macrophages were more prevalent in the virus-high samples. Fibroblasts were abundant in high-virus patient LN3 but also in low-virus patient LN2. plasma cells and airway epithelial cells were particularly prevalent in low-virus patient LN2. However, it should be noted that the tissue section for patient LN2 captured more morphological features than the other patients (i.e. the airway and a portion of smooth muscle), so the abundance of particular cell types in this sample may be driven more by the structural features within the section, than differences in COVID-19 response in this tissue. The other tissues (LN1, LN3 and LN4) were more homogenous within a section, and more similar to one another).

The four patient samples were split into high and low viral levels based on previous work from our group (15, 39). We sought to compare the expression of genes between the two high virus-level and the two low virus-level samples. We identified a total of 115 genes that were upregulated in the high-viral samples, and 2017 genes that were upregulated in the low-viral sample. As expected, visualisation of the expression of all genes together, in the form of an AUC score, indicated highest expression of the high-viral genes in LN1 and LN3 and high expression of the low-viral genes in sample LN2 (**Fig S4c-d**); gene expression was comparatively lower in sample LN4 (**Fig S3c-d**). The high-virus genes are enriched for GO terms including interferon response, antigen processing and chemokine-mediated signalling (**Fig S4f**). In line with the interferon-related role of the upregulated DEGs in the virus-high gene set, 12 of the DEGs are known interferon-related genes (B. Tang, personal communication, 01/08/2021) and these show a clear, qualitative enrichment pattern in LN1 and, in particular, LN3 (**Fig S4e**). Differentially expressed genes (DEGs) between virus-high and virus-low samples were identified using a negative binomial test as implemented in Seurat v4.0.0 (40, 41); GO analysis was performed using clusterProfiler (40) against the org.Hs.eg.db database and a background universe of all expressed genes in the dataset. Resulting DEGs were further filtered by overlapping with genes linked to KEGG pathways hsa05171 (“Coronavirus disease - COVID-19 - Homo sapiens (human)”), hsa04060 (“Cytokine-cytokine receptor interaction - Homo sapiens (human)”), or a curated list of known interferon-related genes (15, 22, 23). Lists of multiple genes were summarised for visualisation by calculating an AUC score (42). Cell type location and proportion per Visium spot was predicted by using label transfer (18, 42) on individual (i.e. unintegrated) samples, based on intermediate cell type annotations from a publicly-available reference dataset (19).

### Image registration analysis of CODEX and Visium data

The Python package *SimpleITK (43)* was used to perform image registration. CODEX images were firstly downscaled to appropriate resolution to match with the resolution of the corresponding Visium histological image. The DAPI channel in the CODEX image was cropped and rotated to have the same capture area and orientation with the Visium histological image and was used as the moving image (query image). Visium histological images were converted to grayscale images to transform the pixel data dimension consistent with the CODEX DAPI channel image and was used as the fixed image (target/reference image). After centralising the two images, the rigid affine transformation was applied for shearing, shifting and scaling the moving image to align with the fixed image in lower resolution as the initial step. Finally, the non-rigid B-spline transformation was applied on affine initialisation to refine the local alignment. The mutual information was used as the evaluation matrix to optimise the parameter for both affine and b-spline transformation.

### Integrating CODEX protein signal to Visium transcriptional data

After registering the CODEX image to Visium histological image, the optimised transformation matrix was then able to convert the cells in CODEX data from the original CODEX spatial coordinates (x, y) to newly mapped spatial coordinates (x’, y’) which are identical to Visium spatial coordinates. With this shared coordinating system, cells in CODEX data then can be searched and grouped by the spatial radius (d=55um, diameter equivalent to Visium spots size) using the transferred spatial coordinates (x’,y’), which created an additional layer of protein expression profiles on top of existing Visium RNA measurements with compatible resolution (Visium spot level). Cell type wise contingency tables were created from binarised CODEX protein marker expression and Visium label transferring results. Chi-squared test was implemented to test the consistency.

### GeoMX data Pre-processing

GeoMX data consists of 49 ROIs in total, 21 belonging to uninfected samples across five patients (N06_062, N06_095, N07_009, N07_034, N08_021) and 28 belonging to COVID-19 infected samples across four patients (LN1, LN2, LN3, LN4). For each ROI, the tissue type annotations (Vessel, T2, Bronchi Epithelium, and Immune) are also available for GeoMX data. GeoMX data has a total of 18704 genes. For the differential gene expression, the 3rd quartile normalised counts provided were further normalised using the edgeR package, which performs library size normalisation. The design matrix included group-level (COVID-19 infected and uninfected) as well as patient-level (9 different patients, 4-COVID-19 and 5-uninfected) information for differential gene expression analysis of all COVID-19 infected vs uninfected samples, for adjusting the differences between nine patients. Whereas for the differential gene expression analysis between high-viral vs low-viral mRNA signal (**Fig S2d**) and COVID-19 infected vs uninfected samples for specific tissue type (Vessel, T2 and Bronchi Epithelium) the design matrix just included the group-level information.

### GeoMX spatial deconvolution

We used the R package *SpatialDecon* (44) to assess the abundance of various cell types for all the ROIs. *SpatialDecon* (44) can estimate cell type abundance for spatially-resolved gene expression studies. The input data required are: reference expression profiles of expected cell types, normalised gene expression matrix and expected background counts of the normalised gene expression at each element calculated by function *derive_GeoMx_background*).

Reference expression profile used here is “SafeTME” (this reference matrix is designed to be biassed towards immune cell types and includes expression of 906 genes to define 18 cell types; some of the known cancer genes are avoided in this profile) we used this reference to identify proportions of different immune cell types across COVID-19 infected and noninfected samples. The overall cell type proportion for all samples is shown in (**Fig S1i**) and the cell types with significant difference in mean are shown in (**Fig S1g**).

### PhenoImager HT cell type and cell community analysis

From the multiplex tissue image, the DAPI channel stained for cell nuclei was used for cell segmentation. By adopting a deep learning model called *stardist* (45) every nuclei from the DAPI channel was transformed into a polygonal cell object. For each cell, mean protein intensity signal was measured and assigned into the cell protein expression level. After several signal preprocessing steps to normalise and remove outlier signal, a standard cell type clustering was applied using a common single-cell processing pipeline, scanpy (33, 45). Based on the panel of six proteins, we were able to identify key immune cell types including NK cells (CD56+), Monocyte/Myeloid (CD15+), naive T cells (CD3+ and CD8+), cytotoxic T-cells (CD8+) and neutrophil (CD66b+) as shown in **Fig S1f**. Among all the clusters detected by the *scanpy* pipeline (Leiden clustering), those cells with very low expression of all the proteins in the panel were classified as unidentified and removed from downstream analysis.

After identifying cell type, we sought to shed light into the distribution of the cell spatial organisations through cell community analysis. For cell community detection, cells are grouped into different clusters of communities using its spatial attributes, which are defined by a K number of nearest neighbouring cells. Clustering of cells with similar spatial neighbourhood patterns (i.e. spatial identity) allows us to identify the composition of cell type for each community.

## Data availability

All of the spatial transcriptomics (Visium and GeoMX WTA), both raw and processed count matrices, will be deposited to ArrayExpress repository (https://www.ebi.ac.uk/arrayexpress/) and raw sequencing data will be available according to human ethics regulations. All other experimental data, including RNAScope, CODEX, PhenoImager HT (imaging data and count data) will be made available in Zenodo.

## Code availability

The code to reproduce analyses and figures presented in this paper is available at https://github.com/BiomedicalMachineLearning/Covid_spatial_multi-omics

## Author contributions statement

Q.N. and A.K. conceived the project and designed the experiment and analysis. A.C.S.F.A., J.S.M.J., K.F.M., C.M.S., P.S.F.G., C.P.B., L.N, F.S.F.G prepared and annotated the clinical samples. T.V., C.U., T.D., S.W., J.S., J.M, A.N, L.P, performed spatial transcriptomics data generation. X.T., L.F.G, M.T., O.M., J.M., T.B., performed data analysis. H.N.L., K.S., G.T.B., F.S.F.G., A.K., and Q.N. performed data interpretation and analysis. All authors contributed to writing the manuscript and approved the final version of the manuscript.

## Acknowledgements

This work was supported by National Health and Medical Research Council (NHMRC) of Australia grants APP2008542 and 1135898 to GTB, 1157741 to A.K, 1140406 to FSFG, 2001514 to QHN, and NHMRC Investigator Grant (GNT2008928) to QHN. The Translational Research Institute is supported by a grant from the Australian Government. We thank the staff at the Institute for Molecular Bioscience for their support in sequencing, the Walter and Eliza Hall Institute (WEHI) histology core, and Enable Medicine.

## Reference

1. Milross, L., Majo, J., Cooper, N., Kaye, P.M., Bayraktar, O., Filby, A. and Fisher, A.J. (2022) Post-mortem lung tissue: the fossil record of the pathophysiology and immunopathology of severe COVID-19. Lancet Respir Med, 10, 95–106.

2. Eyre, D.W., Taylor, D., Purver, M., Chapman, D., Fowler, T., Pouwels, K.B., Walker, A.S. and Peto, T.E.A. (2022) Effect of Covid-19 Vaccination on Transmission of Alpha and Delta Variants. N. Engl. J. Med., 386, 744–756.

3. Nalbandian, A., Sehgal, K., Gupta, A., Madhavan, M.V., McGroder, C., Stevens, J.S., Cook, J.R., Nordvig, A.S., Shalev, D., Sehrawat, T.S., et al. (2021) Post-acute COVID-19 syndrome. Nat. Med., 27, 601–615.

4. McElvaney, O.J., McEvoy, N.L., McElvaney, O.F., Carroll, T.P., Murphy, M.P., Dunlea, D.M., Ní Choileáin, O., Clarke, J., O‘Connor, E., Hogan, G., et al. (2020) Characterization of the Inflammatory Response to Severe COVID-19 Illness. Am. J. Respir. Crit. Care Med., 202, 812–821.

5. Wang, X., Xu, G., Liu, X., Liu, Y., Zhang, S. and Zhang, Z. (2021) Multiomics: unraveling the panoramic landscapes of SARS-CoV-2 infection. Cell. Mol. Immunol., 18, 2313–2324.

6. Biancolella, M., Colona, V.L., Mehrian-Shai, R., Watt, J.L., Luzzatto, L., Novelli, G. and Reichardt, J.K.V. (2022) COVID-19 2022 update: transition of the pandemic to the endemic phase. Hum. Genomics, 16, 19.

7. Knyazev, S., Chhugani, K., Sarwal, V., Ayyala, R., Singh, H., Karthikeyan, S., Deshpande, D., Baykal, P.I., Comarova, Z., Lu, A., et al. (2022) Unlocking capacities of genomics for the COVID-19 response and future pandemics. Nat. Methods, 19, 374–380.

8. Lu, T., Wang, Y. and Guo, T. (2022) Multi-omics in COVID-19: Seeing the unseen but overlooked in the clinic. Cell Rep Med, 3, 100580.

9. Kim, D.-K., Weller, B., Lin, C.-W., Sheykhkarimli, D., Knapp, J.J., Dugied, G., Zanzoni, A., Pons, C., Tofaute, M.J., Maseko, S.B., et al. (2022) A proteome-scale map of the SARS-CoV-2–human contactome. Nat. Biotechnol.

10. Stukalov, A., Girault, V., Grass, V., Karayel, O., Bergant, V., Urban, C., Haas, D.A., Huang, Y., Oubraham, L., Wang, A., et al. (2021) Multilevel proteomics reveals host perturbations by SARS-CoV-2 and SARS-CoV. Nature, 594, 246–252.

11. Delorey, T.M., Ziegler, C.G.K., Heimberg, G., Normand, R., Yang, Y., Segerstolpe, Å., Abbondanza, D., Fleming, S.J., Subramanian, A., Montoro, D.T., et al. (2021) COVID-19 tissue atlases reveal SARS-CoV-2 pathology and cellular targets. Nature, 595, 107–113.

12. Stephenson, E., Reynolds, G., Botting, R.A., Calero-Nieto, F.J., Morgan, M.D., Tuong, Z.K., Bach, K., Sungnak, W., Worlock, K.B., Yoshida, M., et al. (2021) Single-cell multi-omics analysis of the immune response in COVID-19. Nat. Med., 27, 904–916.

13. Li, C.-X., Wheelock, C.E., Sköld, C.M. and Wheelock, Å.M. (2018) Integration of multi-omics datasets enables molecular classification of COPD. Eur. Respir. J., 51.

14. Rendeiro, A.F., Ravichandran, H., Bram, Y., Chandar, V., Kim, J., Meydan, C., Park, J., Foox, J., Hether, T., Warren, S., et al. (2021) The spatial landscape of lung pathology during COVID-19 progression. Nature, 593, 564–569.

15. Kulasinghe, A., Tan, C.W., Ribeiro Dos Santos Miggiolaro, A.F., Monkman, J., SadeghiRad, H., Bhuva, D.D., Motta Junior, J. da S., Busatta Vaz de Paula, C., Nagashima, S., Baena, C.P., et al. (2022) Profiling of lung SARS-CoV-2 and influenza virus infection dissects virus-specific host responses and gene signatures. Eur. Respir. J., 59.

16. Lewis, S.M., Asselin-Labat, M.-L., Nguyen, Q., Berthelet, J., Tan, X., Wimmer, V.C., Merino, D., Rogers, K.L. and Naik, S.H. (2021) Spatial omics and multiplexed imaging to explore cancer biology. Nat. Methods, 18, 997–1012.

17. Tran, M., Yoon, S., Teoh, M., Andersen, S., Lam, P.Y., Purdue, B.W., Raghubar, A., Hanson, S.J., Devitt, K., Jones, K., et al. (2022) A robust experimental and computational analysis framework at multiple resolutions, modalities and coverages. Front. Immunol., 13, 911873.

18. Stuart, T., Butler, A., Hoffman, P., Hafemeister, C., Papalexi, E., Mauck, W.M., Hao, Y., Stoeckius, M., Smibert, P. and Satija, R. (2019) Comprehensive Integration of Single-Cell Data. Cell, 177.

19. Melms, J.C., Biermann, J., Huang, H., Wang, Y., Nair, A., Tagore, S., Katsyv, I., Rendeiro, A.F., Amin, A.D., Schapiro, D., et al. (2021) A molecular single-cell lung atlas of lethal COVID-19. Nature, 595.

20. Reyes, L., Sanchez-Garcia, M.A., Morrison, T., Howden, A.J.M., Watts, E.R., Arienti, S., Sadiku, P., Coelho, P., Mirchandani, A.S., Zhang, A., et al. (2021) A type I IFN, prothrombotic hyperinflammatory neutrophil signature is distinct for COVID-19 ARDS. Wellcome Open Research, 6.

21. Montazersaheb, S., Hosseiniyan Khatibi, S.M., Hejazi, M.S., Tarhriz, V., Farjami, A., Ghasemian Sorbeni, F., Farahzadi, R. and Ghasemnejad, T. (2022) COVID-19 infection: an overview on cytokine storm and related interventions. Virol. J., 19, 92.

22. Thoutam, A., Breitzig, M., Lockey, R. and Kolliputi, N. (2020) Coronavirus: a shift in focus away from IFN response and towards other inflammatory targets. J. Cell Commun. Signal.

23. Amanat, F. and Krammer, F. (2020) SARS-CoV-2 Vaccines: Status Report. Immunity, 52, 583–589.

24. Farber, J.M. (1990) A macrophage mRNA selectively induced by gamma-interferon encodes a member of the platelet factor 4 family of cytokines. Proc. Natl. Acad. Sci. U. S. A., 87, 5238–5242.

25. Callahan, V., Hawks, S., Crawford, M.A., Lehman, C.W., Morrison, H.A., Ivester, H.M., Akhrymuk, I., Boghdeh, N., Flor, R., Finkielstein, C.V., et al. (2021) The Pro-Inflammatory Chemokines CXCL9, CXCL10 and CXCL11 Are Upregulated Following SARS-CoV-2 Infection in an AKT-Dependent Manner. Viruses, 13.

26. Li, C.-X., Gao, J., Zhang, Z., Chen, L., Li, X., Zhou, M. and Wheelock, Å.M. (2022) Multiomics integration-based molecular characterizations of COVID-19. Brief. Bioinform., 23.

27. Barh, D., Tiwari, S., Weener, M.E., Azevedo, V., Góes-Neto, A., Gromiha, M.M. and Ghosh, P. (2020) Multi-omics-based identification of SARS-CoV-2 infection biology and candidate drugs against COVID-19. Comput. Biol. Med., 126, 104051.

28. Su, A., Lee, H., Tan, X., Suarez, C.J., Andor, N., Nguyen, Q. and Ji, H.P. (2022) A deep learning model for molecular label transfer that enables cancer cell identification from histopathology images. NPJ Precis Oncol, 6, 14.

29. Grant, R.A., Morales-Nebreda, L., Markov, N.S., Swaminathan, S., Querrey, M., Guzman, E.R., Abbott, D.A., Donnelly, H.K., Donayre, A., Goldberg, I.A., et al. (2021) Circuits between infected macrophages and T cells in SARS-CoV-2 pneumonia. Nature, 590, 635–641.

30. Jafarzadeh, A., Chauhan, P., Saha, B., Jafarzadeh, S. and Nemati, M. (2020) Contribution of monocytes and macrophages to the local tissue inflammation and cytokine storm in COVID-19: Lessons from SARS and MERS, and potential therapeutic interventions. Life Sciences, 257, 118102.

31. Black, S., Phillips, D., Hickey, J.W., Kennedy-Darling, J., Venkataraaman, V.G., Samusik, N., Goltsev, Y., Schürch, C.M. and Nolan, G.P. (2021) CODEX multiplexed tissue imaging with DNA-conjugated antibodies. Nat. Protoc., 16, 3802–3835.

32. Bannon, D., Moen, E., Schwartz, M., Borba, E., Kudo, T., Greenwald, N., Vijayakumar, V., Chang, B., Pao, E., Osterman, E., et al. (2021) DeepCell Kiosk: scaling deep learning-enabled cellular image analysis with Kubernetes. Nat. Methods, 18, 43–45.

33. Wolf, F.A., Angerer, P. and Theis, F.J. (2018) SCANPY: large-scale single-cell gene expression data analysis. Genome Biol., 19, 15.

34. McCarthy, D.J., Campbell, K.R., Lun, A.T.L. and Wills, Q.F. (2017) Scater: pre-processing, quality control, normalization and visualization of single-cell RNA-seq data in R. Bioinformatics, 33, 1179–1186.

35. Chen, Y., Lun, A.T.L. and Smyth, G.K. (2016) From reads to genes to pathways: differential expression analysis of RNA-Seq experiments using Rsubread and the edgeR quasi-likelihood pipeline. F1000Res., 5, 1438.

36. Lun, A.T.L., McCarthy, D.J. and Marioni, J.C. (2016) A step-by-step workflow for low-level analysis of single-cell RNA-seq data with Bioconductor. F1000Res., 5, 2122.

37. Butler, A., Hoffman, P., Smibert, P., Papalexi, E. and Satija, R. (2018) Integrating single-cell transcriptomic data across different conditions, technologies, and species. Nat. Biotechnol., 36, 411–420.

38. Zappia, L. and Oshlack, A. (2018) Clustering trees: a visualization for evaluating clusterings at multiple resolutions. GigaScience, 7.

39. Shojaei, M., Shamshirian, A., Monkman, J., Grice, L., Tran, M., Tan, C.W., Rossi, G.R., McCulloch, T.R., Nalos, M., Chew, K.Y., et al. (2021) IFI27 transcription is an early predictor for COVID-19 outcomes; a multi-cohort observational study. medRxiv, 10.1101/2021.10.29.21265555.

40. Wu, T., Hu, E., Xu, S., Chen, M., Guo, P., Dai, Z., Feng, T., Zhou, L., Tang, W., Zhan, L., et al. (2021) clusterProfiler 4.0: A universal enrichment tool for interpreting omics data. Innovation (Camb), 2, 100141.

41. Integrated analysis of multimodal single-cell data (2021) Cell, 184, 3573–3587.e29.

42. Aibar, S., González-Blas, C.B., Moerman, T., Huynh-Thu, V.A., Imrichova, H., Hulselmans, G., Rambow, F., Marine, J.-C., Geurts, P., Aerts, J., et al. (2017) SCENIC: single-cell regulatory network inference and clustering. Nat. Methods, 14, 1083–1086.

43. Beare, R., Lowekamp, B. and Yaniv, Z. (2018) Image Segmentation, Registration and Characterization in R with SimpleITK. J. Stat. Softw., 86.

44. Danaher, P., Kim, Y., Nelson, B., Griswold, M., Yang, Z., Piazza, E. and Beechem, J.M. (2022) Advances in mixed cell deconvolution enable quantification of cell types in spatial transcriptomic data. Nat. Commun., 13, 1–13.

45. Stevens, M., Nanou, A., Terstappen, L.W.M.M., Driemel, C., Stoecklein, N.H. and Coumans, F.A.W. (2022) StarDist Image Segmentation Improves Circulating Tumor Cell Detection. Cancers, 14.

