## Supplemental Figures for "A robust Platform for Integrative Spatial Multi-omics Analysis to Map Immune Responses to SARS-CoV-2 infection in Lung Tissues"

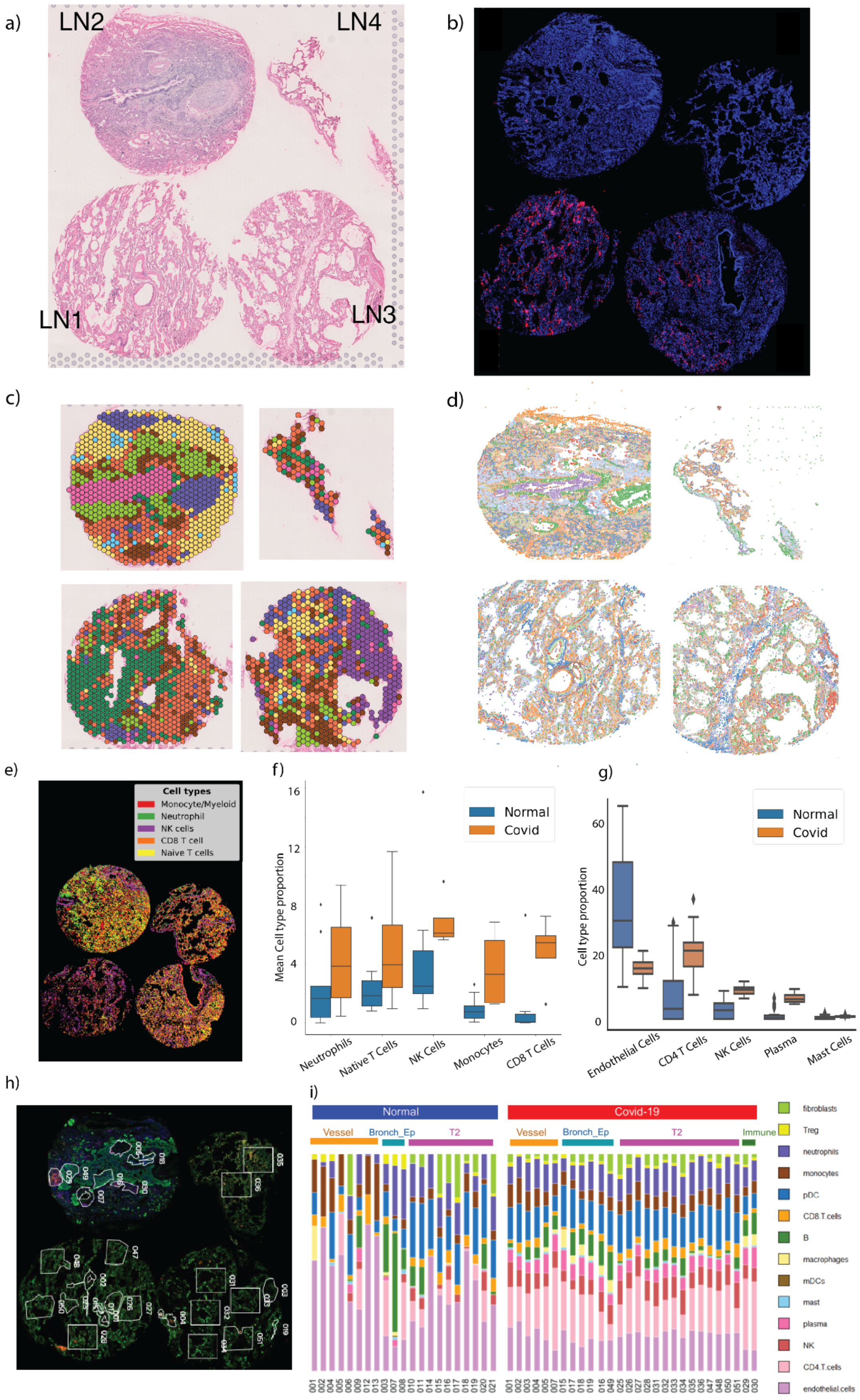

a)
LN2
LN4
LN1
LN3
b)
c)
d)
e)
Cell types
Monocyte/Myeloid
Neutrophil
NK cells
CD8 T cell
Naive T cells
f)
Normal
Covid
Mean Cell type proportion
Neutrophils
Native T Cells
NK Cells
Monocytes
CD8 T Cells
g)
Normal
Covid
Cell type proportion
Endothelial Cells
CD4 T Cells
NK Cells
Plasma
Mast Cells
h)
i)
Normal
Covid-19
Vessel
Bronch_Ep
T2
Immune
fibroblasts
Treg
neutrophils
monocytes
pDC
CD8.T.cells
B
macrophages
mDCs
mast
plasma
NK
CD4.T.cells
endothelial.cells

**Supplementary Figure 1. Comparison of cell type composition and distribution defined by each of the five spatial technologies applied across infected tissues in four COVID-19 patients.**

(a) H&E stain of lung biopsies from four patients (LN1-LN4). LN1 and LN3 were defined as having a high viral mRNA signal, while LN2 and LN4 had a low viral signal, as defined by RNAScope (Panel b). The tissue image shown was collected as part of the Visium protocol (Panel c) but adjacent sections from the same tissue block were used for generating Phenoimager HT data. Additional sections from the same block, but further away were compared to the Phenoimager HT section, were used for PhenoImager HT, RNAScope and GeoMx data generation.
(b) RNAScope detection of SARS-CoV-2 viral mRNA in lung tissue sections. Red colour indicates positive detection of the viral mRNA.
(c) Spot clustering results from Visium data. Each colour represents a cluster; cluster colours are consistent across tissues.
(d) Independent clustering of different cell types using Phenoimager HT data. Each tissue was clustered separately so colours are not consistent across tissues or data modalities. Phenoimager HT technology was applied to tissue sections adjacent to those used for Visium data generation; sections used for PhenoImager HT, RNAScope and GeoMx were taken from further away in the block.
(e) PhenoImager HT cell type classification using combined expression of six protein markers.
(f) Comparison of cell type proportions between COVID-19-infected and uninfected samples using PhenoImager HT data. Y-axis shows the mean cell type percentage between replicates (four COVID-19 and nine uninfected samples).
(g) Comparison of cell type proportions between COVID-19-infected and uninfected samples using GeoMX data. Cell types are arranged along the x-axis in the order of statistical significance of the mean difference between conditions.
(h) GeoMX COVID-19 tissues with 49 ROIs indicated as white polygons.
(i) Cell type composition across ROIs from uninfected and COVID-19-infected samples in GeoMX data. ROIs from infected samples contain CD4 T cells, NK Cells and plasma cells in abundance compared to healthy controls, while healthy ROIs show enrichment for endothelial cells. ROI: region of interest.

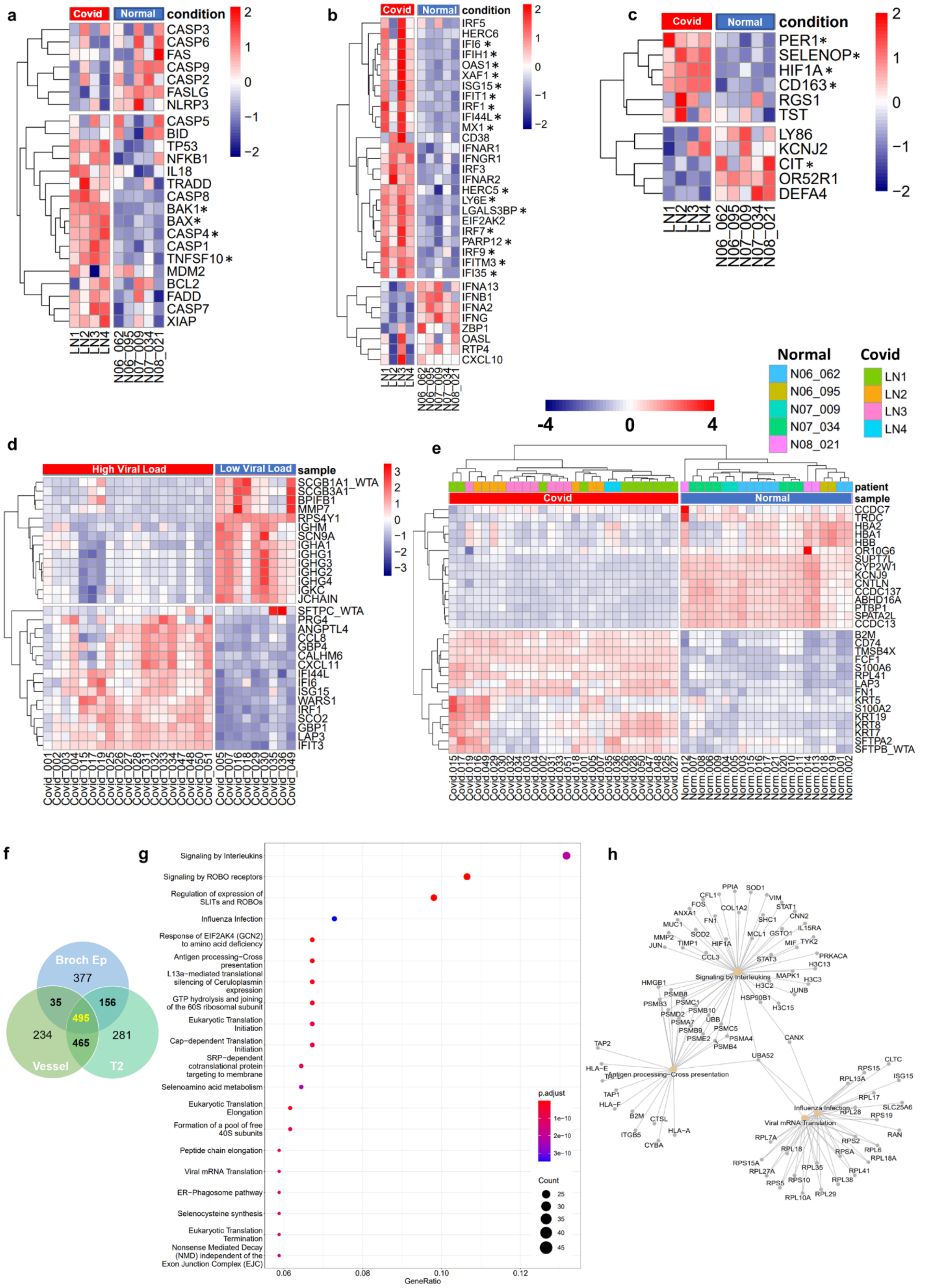

a
Covid
Normal
condition
CASP3
CASP6
FAS
CASP9
CASP2
FASLG
NLRP3
CASP5
BID
TP53
NFKB1
IL18
TRADD
CASP8
BAK1*
BAX*
CASP4*
CASP1
TNFSF10*
MDM2
BCL2
FADD
CASP7
XIAP
LN1
LN2
LN3
LN4
N06_062
N06_095
N07_009
N07_034
N08_021
b
IRF5
HERC6
IFI6*
IFIH1*
OAS1*
XAF1*
ISG15*
IFIT1*
IRF1*
IFI44L*
MX1*
CD38
IFNAR1
IFNGR1
IRF3
IFNAR2
HERC5*
LY6E*
LGALS3BP*
EIF2AK2
IRF7*
PARP12*
IRF9*
IFITM3*
IFI35*
IFNA13
IFNB1
IFNA2
IFNG
ZBP1
OASL
RTP4
CXCL10
c
PER1*
SELENOP*
HIF1A*
CD163*
RGS1
TST
LY86
KCNJ2
CIT*
OR52R1
DEFA4
Normal
N06_062
N06_095
N07_009
N07_034
N08_021
Covid
LN1
LN2
LN3
LN4
-4
0
4
d
High Viral Load
Low Viral Load
sample
SCGB1A1_WTA
SCGB3A1
BPIFB1
MMP7
RPS4Y1
IGHM
SCN9A
IGHA1
IGHG1
IGHG3
IGHG2
IGHG4
IGKC
JCHAIN
SFTPC_WTA
PRG4
ANGPTL4
CCL8
GBP4
CALHM6
CXCL11
IFI44L
IFI6
ISG15
WARS1
IRF1
SCO2
GBP1
LAP3
IFIT3
e
patient
sample
Covid
Normal
CCDC7
TRDC
HBA2
HBA1
HBB
OR10G6
SUPT7L
CYP2W1
KCNJ9
CNTLN
CCDC137
ABHD16A
PTBP1
SPATA2L
CCDC13
B2M
CD74
TMSB4X
FCF1
S100A6
RPL41
LAP3
FN1
KRT5
S100A2
KRT19
KRT8
KRT7
SFTPA2
SFTPB_WTA
f
Broch Ep
377
35
156
495
234
465
281
Vessel
T2
g
Signaling by Interleukins
Signaling by ROBO receptors
Regulation of expression of SLITs and ROBOs
Influenza Infection
Response of EIF2AK4 (GCN2) to amino acid deficiency
Antigen processing−Cross presentation
L13a−mediated translational silencing of Ceruloplasmin expression
GTP hydrolysis and joining of the 60S ribosomal subunit
Eukaryotic Translation Initiation
Cap−dependent Translation Initiation
SRP−dependent cotranslational protein targeting to membrane
Selenoamino acid metabolism
Eukaryotic Translation Elongation
Formation of a pool of free 40S subunits
Peptide chain elongation
Viral mRNA Translation
ER−Phagosome pathway
Selenocysteine synthesis
Eukaryotic Translation Termination
Nonsense Mediated Decay (NMD) independent of the Exon Junction Complex (EJC)
p.adjust
1e−10
2e−10
3e−10
Count
25
30
35
40
45
0.06
0.08
0.10
0.12
GeneRatio
h
Signaling by Interleukins
Antigen processing−Cross presentation
Influenza Infection
Viral mRNA Translation

**Supplementary Figure 2. Nanostring GeoMX transcriptome-wide expression analysis of Covid infected vs non-infected samples and comparisons of high- vs low- virus load across the three sites.**

**a**-**c**) Comparing gene expression between covid infected and non-covid infected samples for previously known or potential markers for the (**a**) Apoptosis responses (**b**) Interferons responses and (**c**) Prognostic markers. Log Counts Per Million (logCPM) transformed values (normalised gene expression counts by edgeR cpm function) were grouped by patients and averaged to look at the differences between covid and normal. Curated gene lists for the above three pathways received from pathologists were used for comparison. (* indicates the genes differentially expressed between Covid and non-Covid infections).

**d-e**) Normalised gene expression counts (edgeR cpm) were used. Genes with FDR <= 0.05 and absolute of log2 fold change >= 1 were categorised as differential genes. (**d**) Top 15 genes upregulated in high virus loading and low virus loading samples sorted by logFC are shown. (**e**) Top 15 genes upregulated in covid and non-covid infected samples sorted by logFC are shown.

**g**) Reactome Pathway Database [1] was used to perform pathway analysis of 495 genes upregulated in covid samples. These genes are consistently differentially expressed in covid and non-covid infected samples across different tissue types (Vesel, Bronchi Epithelium and T2). 481 genes here also overlap with the DE gene list of covid vs non-covid infected samples, further indicating the biological importance of these genes. Top 20 pathways are shown.

**h**) Cnetplot of four pathways (amongst the top 20 pathways) associated with Covid infection is shown.

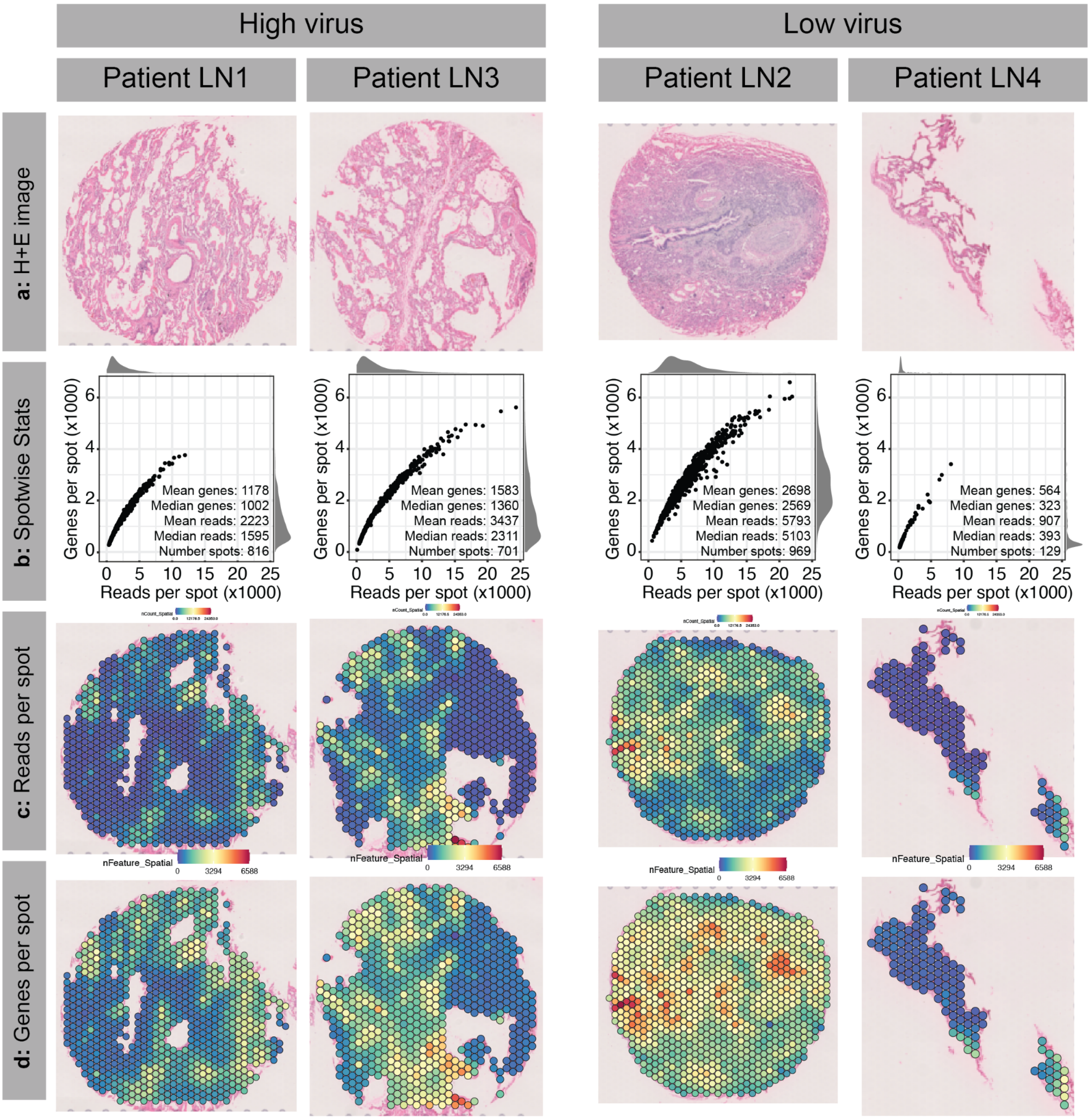


**Supplementary Figure 3. Histological tissue images and quality metrics in Visium spatial transcriptomics data.**

**a**) H&E stained images showing the four SARS-CoV-2 lung samples used for Visium spatial transcriptomics. Samples are categorised as high-viral or low-viral load based on staining to target the SARS-CoV-2 spike mRNA using RNAScope.

**b**) Summary statistics for each sample. Scatter plots show the relationship between captured genes (y-axis) and reads per spot (x-axis) per spot. Other quality metrics are listed for each sample.

**c-d**) Data quality across the samples as indicated by the number of reads (**c**) or genes (**d**) per spatial spot

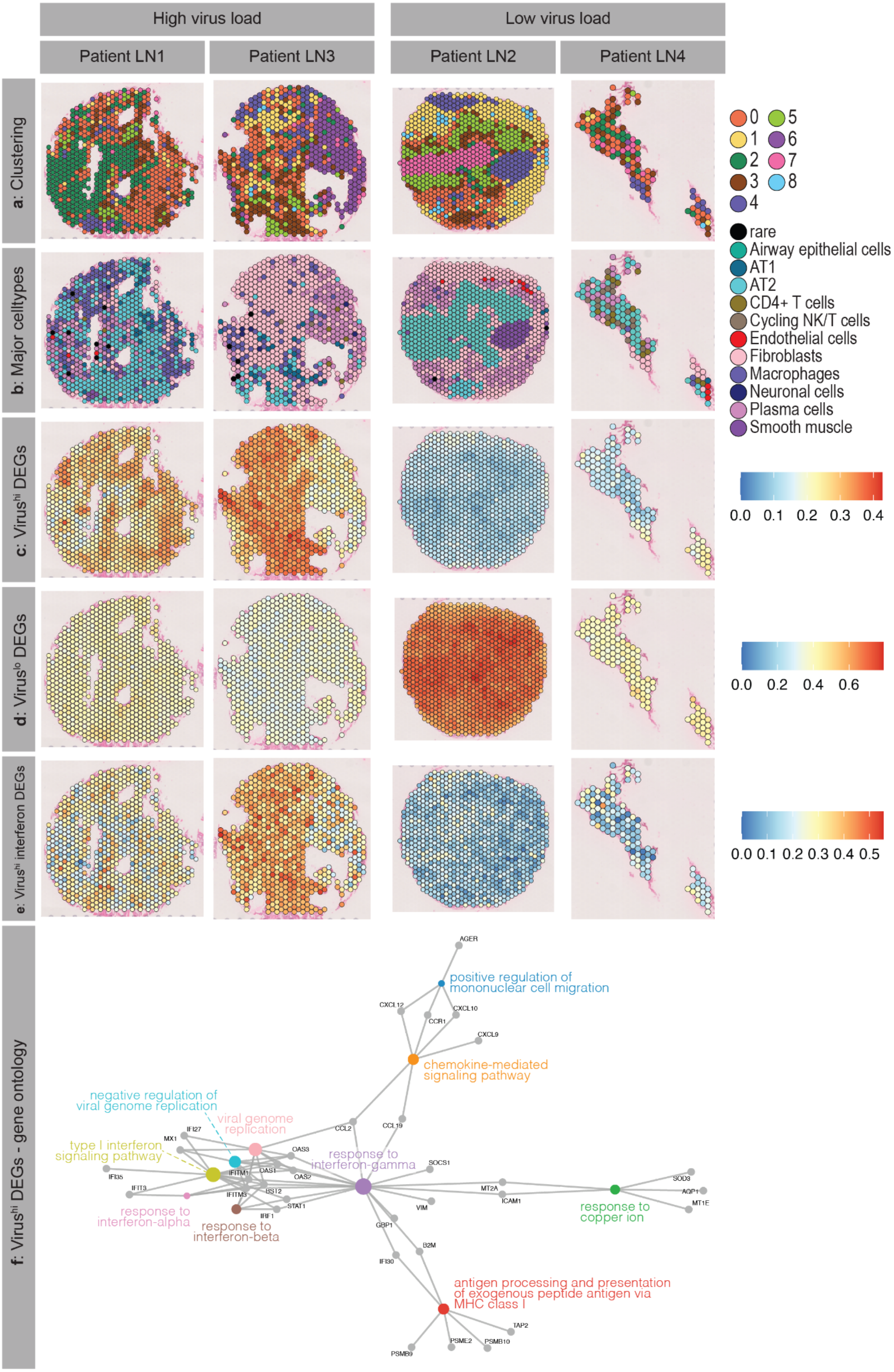

High virus load
Low virus load
Patient LN1
Patient LN3
Patient LN2
Patient LN4
a: Clustering
b: Major celltypes
c: Virus^hi DEGs
d: Virus^lo DEGs
e: Virus^hi interferon DEGs
f: Virus^hi DEGs - gene ontology
0 1 2 3 4 5 6 7 8
rare
Airway epithelial cells
AT1
AT2
CD4+ T cells
Cycling NK/T cells
Endothelial cells
Fibroblasts
Macrophages
Neuronal cells
Plasma cells
Smooth muscle
0.0 0.1 0.2 0.3 0.4
0.0 0.2 0.4 0.6
0.0 0.1 0.2 0.3 0.4 0.5
AGER
positive regulation of mononuclear cell migration
CXCL12
CXCL10
CCR1
CXCL9
chemokine-mediated signaling pathway
negative regulation of viral genome replication
viral genome replication
type I interferon signaling pathway
IFI27
MX1
CCL2
CCL19
OAS3
response to interferon-gamma
SOCS1
IFI35
IFITM1
OAS1
OAS2
IFIT3
IFITM3
BST2
STAT1
IRF1
MT2A
ICAM1
VIM
SOD3
AQP1
MT1E
response to copper ion
response to interferon-alpha
response to interferon-beta
GBP1
B2M
IFI30
antigen processing and presentation of exogenous peptide antigen via MHC class I
TAP2
PSME2
PSMB10
PSMB9

**Supplementary Figure 4. Cell types and gene expression in high- and low-viral infected Visium samples**

**a**) Results of clustering on integrated data (clustering resolution of 0.6).

**b**) Consensus results of label transfer to annotate spots. Label transfer was performed using the "Intermediate" annotation from Melms. et al. [2] run on each sample individually in turn. Spots are coloured by their dominant predicted cell type. Cell types present in four or fewer spots across the entire dataset were renamed as "rare" (black).

**c-d**) Expression of genes differentially expressed between high and low viral samples, split into genes upregulated (**c**) or downregulated (**d**) in high-virus samples compared to low-virus samples. Expression is represented as AUC scores comparing each gene list to the entire transcriptome. AUC: area under the curve

**e**) AUC scores showing expression of genes upregulated in high-virus samples that are also members of a curated list of known interferon genes.

**f**) Gene ontology network diagram of gene ontology enrichment calculated from genes upregulated in high-virus samples. No significant GO terms were detected from the list of genes enriched in low-virus samples. GO: Gene ontology

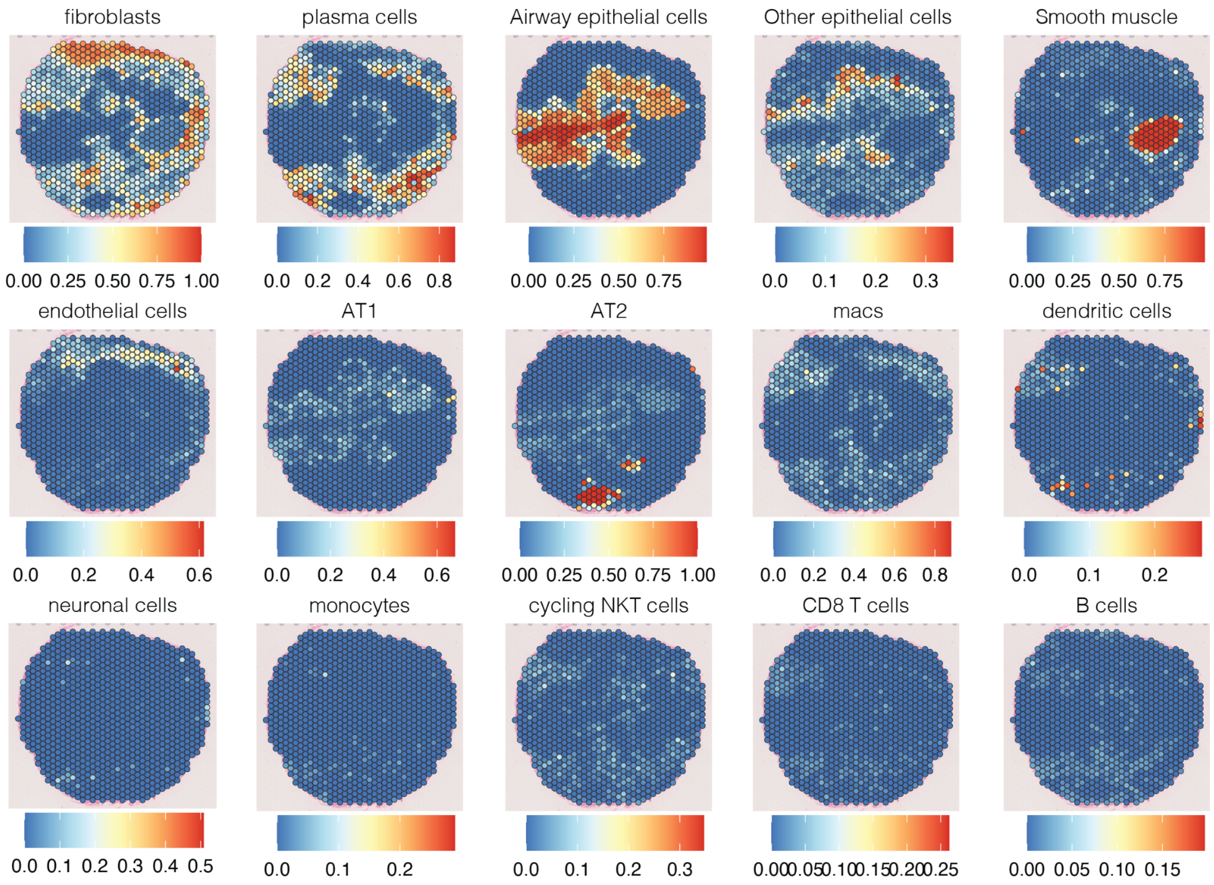


**Supplementary Figure 5. Label transfer scores for all cell types predictions shown in B plotted to the tissue from a COVID-19 infected sample (LN2 patient).**

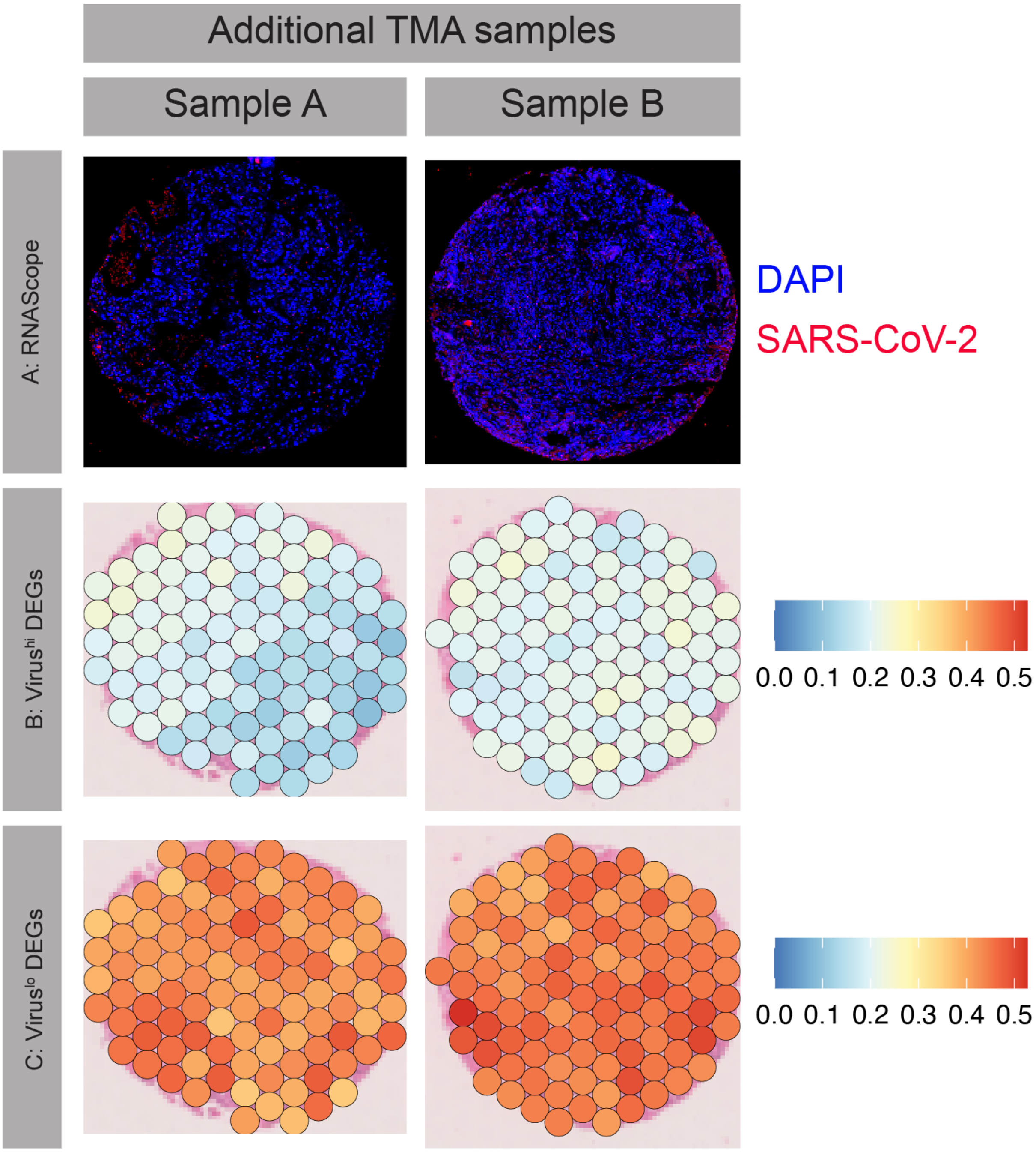


**Supplementary Figure 6. Visualisation of expression of two key gene signatures in an independent COVID-19 infected sample**

**a)** RNAScope detection of SARS-CoV-2 viral mRNA in adjacent lung tissue sections from two technical replicates of one additional COVID-19 patient in a separate cohort. The red colour indicates positive detection of the viral mRNA. Both samples were categorised as having low COVID-19 viral loads.

**b-c**) AUC gene set activity scores of two gene signatures found upregulated genes (**b**) in high-virus samples (**b**) or upregulated genes in low-virus samples (**c**). The two signatures were found in the analysis of the four core Visium COVID-19 samples and validated in an independent sample (with two technical replicates from two adjacent sections).

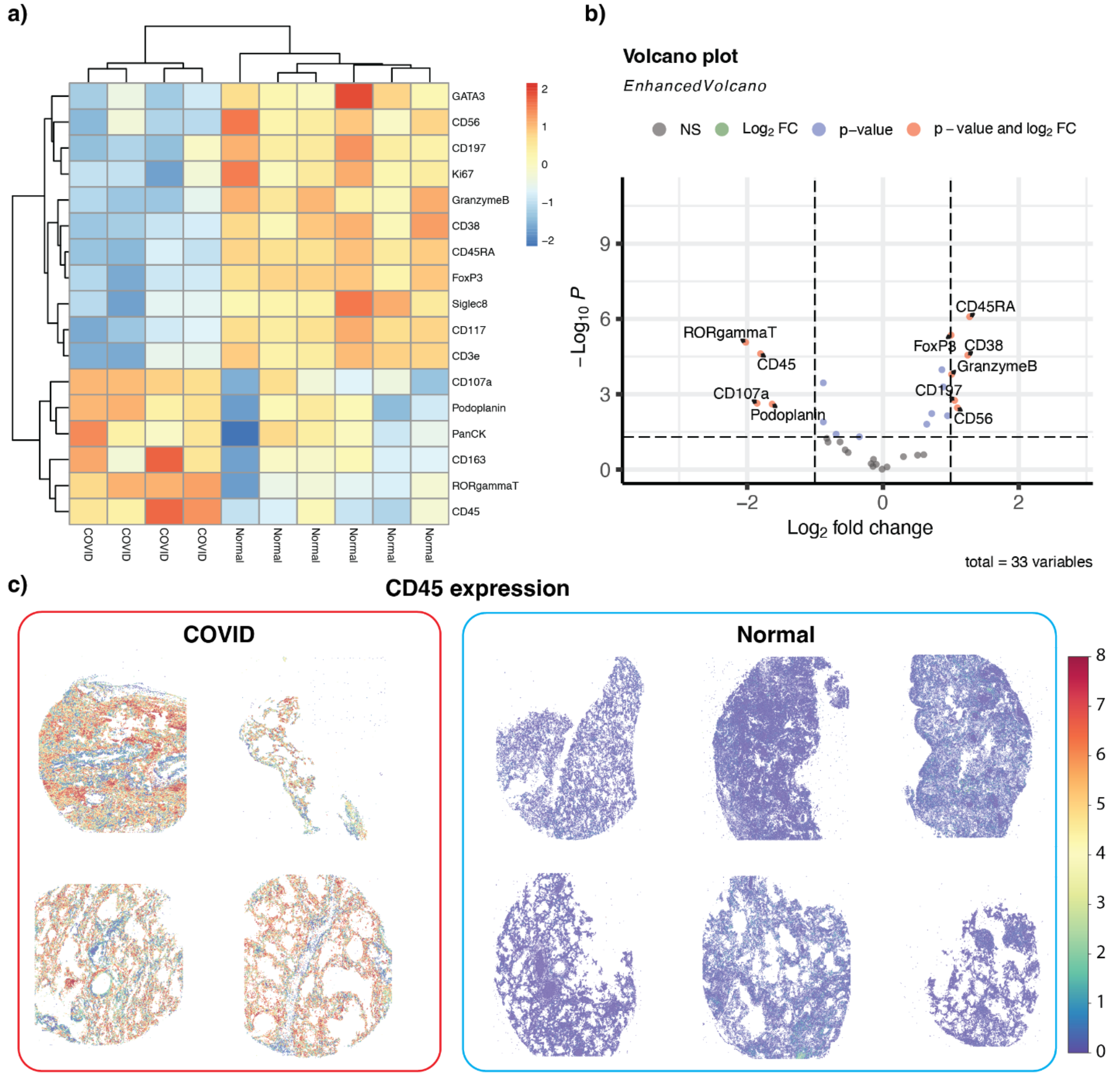


**Supplementary Figure 7. Differential expression analysis of Phenoimager HT data for COVID-19 infected vs non-infected samples using Phenoimager HT data**

**a-b**) Heatmap and volcano plots show the 17 significant differentially expressed protein markers (a) and all 33 markers (b).

**c**) Spatial expression of immune marker CD45 in COVID-19 infected and normal lung tissue clearly shows the strong immune responses in the COVID-19 infected samples.

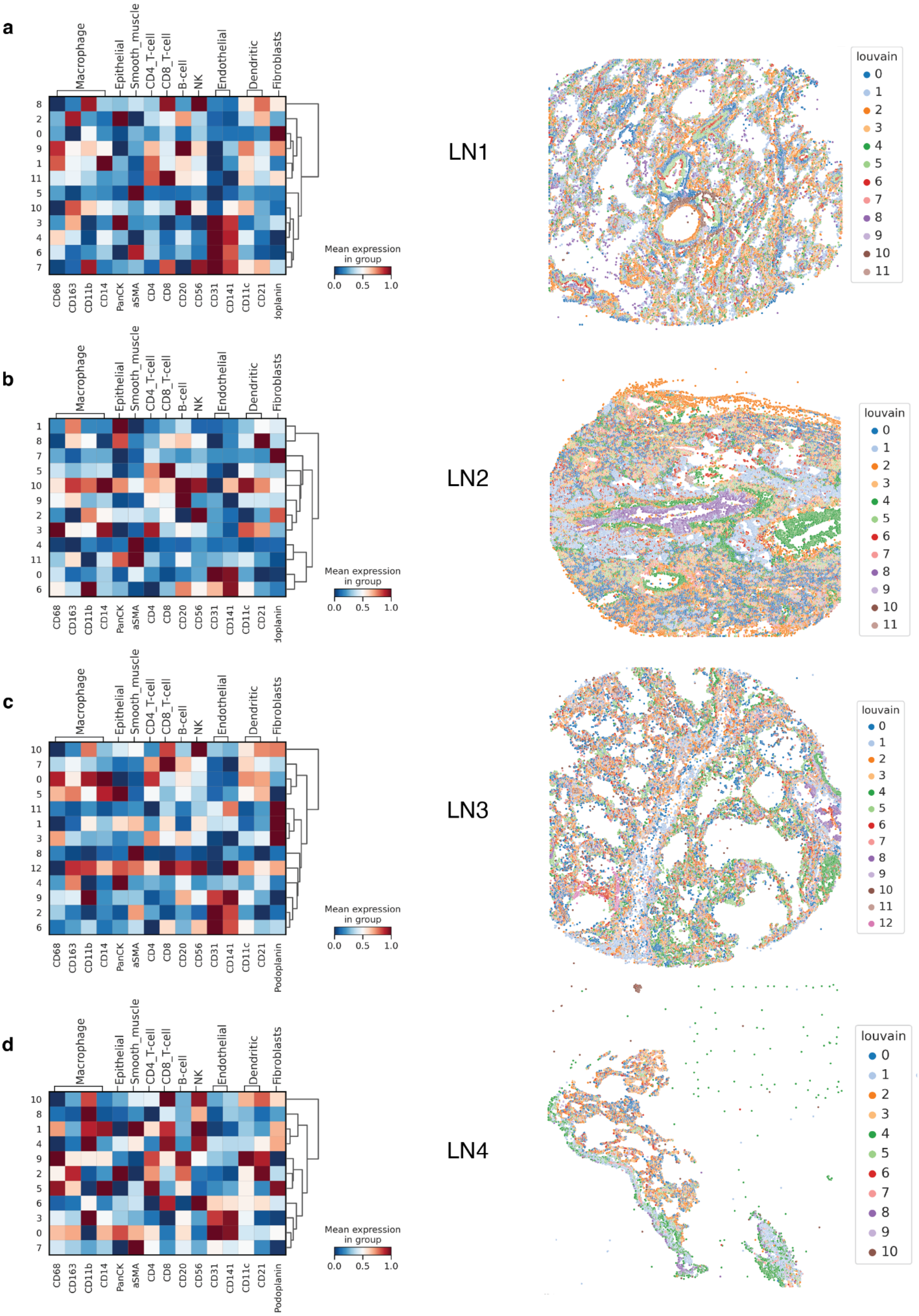
a
Macrophage
Epithelial
Smooth_muscle
CD4_T-cell
CD8_T-cell
B-cell
NK
Endothelial
Dendritic
Fibroblasts
8 2 0 9 1 11 5 10 3 4 6 7
CD68 CD163 CD11b CD14 PanCK aSMA CD4 CD8 CD20 CD56 CD31 CD141 CD11c CD21 doplanin
Mean expression in group
0.0 0.5 1.0
LN1
louvain
0 1 2 3 4 5 6 7 8 9 10 11
b
Macrophage
Epithelial
Smooth_muscle
CD4_T-cell
CD8_T-cell
B-cell
NK
Endothelial
Dendritic
Fibroblasts
1 8 7 5 10 9 2 3 4 11 0 6
CD68 CD163 CD11b CD14 PanCK aSMA CD4 CD8 CD20 CD56 CD31 CD141 CD11c CD21 doplanin
Mean expression in group
0.0 0.5 1.0
LN2
louvain
0 1 2 3 4 5 6 7 8 9 10 11
c
Macrophage
Epithelial
Smooth_muscle
CD4_T-cell
CD8_T-cell
B-cell
NK
Endothelial
Dendritic
Fibroblasts
10 7 0 5 11 1 3 8 12 4 9 2 6
CD68 CD163 CD11b CD14 PanCK aSMA CD4 CD8 CD20 CD56 CD31 CD141 CD11c CD21 Podoplanin
Mean expression in group
0.0 0.5 1.0
LN3
louvain
0 1 2 3 4 5 6 7 8 9 10 11 12
d
Macrophage
Epithelial
Smooth_muscle
CD4_T-cell
CD8_T-cell
B-cell
NK
Endothelial
Dendritic
Fibroblasts
10 8 1 4 9 2 5 6 3 0 7
CD68 CD163 CD11b CD14 PanCK aSMA CD4 CD8 CD20 CD56 CD31 CD141 CD11c CD21 Podoplanin
Mean expression in group
0.0 0.5 1.0
LN4
louvain
0 1 2 3 4 5 6 7 8 9 10

**Supplementary Figure 8. Correspondence between cell type clustering and known markers.**

**a-d**) Left panel, heatmaps show the mean expression of each marker (X-axis) in different clusters (Y-axis). Markers are ordered by cell types (columns). Right panel, Louvain clustering for each sample based on Phenoimager HT data; each dot represents a cell, and each colour shows a cell cluster. The heatmaps show a common trend in all four samples that multiple clusters have colocalisation of  macrophages colocalise with CD8T-cell, Dendritic cells and NK cells.

### Reference


1. Fabregat, A. *et al.* Reactome pathway analysis: a high-performance in-memory approach. *BMC Bioinformatics* **18**, 1–9 (2017).
2. Melms, J. C. *et al.* A molecular single-cell lung atlas of lethal COVID-19. *Nature* **595**, 114–119 (2021).
